# Discovery and Biosynthesis of Nyuzenamides D and E by Genome Mining in *Streptomyces hygroscopicus*

**DOI:** 10.1101/2024.12.20.629799

**Authors:** Kah Yean Lum, Rohitesh Kumar, Zhijie Yang, Luciano D. O. Souza, Scott A. Jarmusch, José M. A. Moreira, Jens Preben Morth, Ling Ding

**Affiliations:** Department of Biotechnology and Biomedicine, Technical University of Denmark, Søltofts Plads 221, DK-2800 Kgs. Lyngby, Denmark; Sino-Danish Center for Education and Research (SDC), Aarhus University, Dalgas Avenue 4, Building 3410, 8000 Aarhus C, Denmark; Department of Drug Design and Pharmacology, Faculty of Health and Medical Sciences, University of Copenhagen, Universitetsparken 2, 2100 Copenhagen, Denmark

## Abstract

Nyuzenamides belong to the class of bioactive cinnamoyl moiety containing non-ribosomal peptides (NRPs). However, their biosynthetic gene cluster (BGC) remains unconfirmed. Genome-mining revealed a putative *nyu* BGC in *Streptomyces hygroscopicus* DSM 40578. Nyuzenamides D (**1**) and E (**2**) were subsequently isolated, and the structures were elucidated by detailed NMR spectroscopic and MS spectrometric data analyses. The absolute configuration of **1** was determined by a single-crystal X-ray diffraction study. Through retro-biosynthesis and CRISPR-genome editing, the non-ribosomal peptide synthetase biosynthetic gene cluster for nyuzenamides was confirmed. Our discovery enriches the diversity of cinnamoyl-containing nonribosomal peptides and validates the biosynthetic gene clusters for future genome-mining research.

## INTRODUCTION

Specialized metabolites produced by the filamentous Gram-positive actinomycetes account for nearly two-thirds of all naturally occurring antibiotics, with the genus *Streptomyces* being the most prolific source.^1^ The major classes of antibiotics produced by *Streptomyces* include peptides, polyenes, aminoglycosides, *b*-lactams, macrolides, ansamycins, to name a few.^2^ Nonribosomal peptides (NRPs) represented by the antibiotic vancomycin, are important group of specialized metabolites synthesized by nonribosomal peptide synthetases (NRPSs). Among them, cinnamoyl-containing NRPs are a small family of microbial natural products and often exhibit diverse bioactivities. Examples include pepticinnamin E, which acts as a farnesyl transferase inhibitor;^3^ atratumycin^4^ and atrovimycin,^5^ both of which are active against *Mycobacterium tuberculosis*; mohangamides A, which inhibit *Candida albicans* isocitrate lyase;^6^ WS9326D, which inhibits *Brugia malayi* asparaginyl-tRNA synthetase;^7^ and coprisamides A and B exhibit activity in inducing quinone reductase.^8^

Nyuzenamides constitute a small family of NRPs containing a cinnamoyl moiety (**Chart 1**). The initial members, nyuzenamides A and B, were isolated from the marine-derived *Streptomyces* sp. N11-34.^9^ These compounds demonstrated inhibitory activities against human and plant pathogenic filamentous fungi, and were cytotoxic against P388 murine leukemia cells.^9^ Subsequently, nyuzenamide C, the third member of the family featuring an EPCA acyl chain, was reported from a *Streptomyces* sp. DM14 isolated from riverine sediment.^10^ Nyuzenamide C did not show cytotoxicity against a panel of bacteria, fungi, and cancer cell lines. However, it exhibited antiangiogenic activity in human umbilical vein endothelial cells.^10^

*Streptomyces hygroscopicus*^11^ can be considered a prolific producer of bioactive secondary metabolites, with approximately 180 compounds described.^12^ Examples are immunosuppressant rapamycin, cytotoxic pterocidin, antifungal validamycin, antibacterial hygromycins, and herbicidal herbimycin.^2^ Additionally, plant growth promoters pteridic acids A and B, were also isolated from this species.^13^ In this study, genome-mining of *S. hygroscopicus* subsp. *hygroscopicus* DSM 40578 revealed 42 biosynthetic gene clusters (BGC), including one undescribed nonribosomal peptide synthetase (NRPS) with similarity to the skyllamycin BGC (**Table S19**). Analysis of the extract from maltose-yeast extract-malt extract (MYM) agar culture by HPLC-HRMS, along with searches in different databases,^14,15^ revealed the presence of previously undescribed NRPs potentially produced by the *nyu* BGC. Here, we describe the isolation, structure elucidation, and biological activities of nyuzenamides D (**1**) and E (**2**). Moreover, we report the biosynthetic gene cluster of nyuzenamides which was confirmed through CRISPR genome editing.

**Chart 1.**
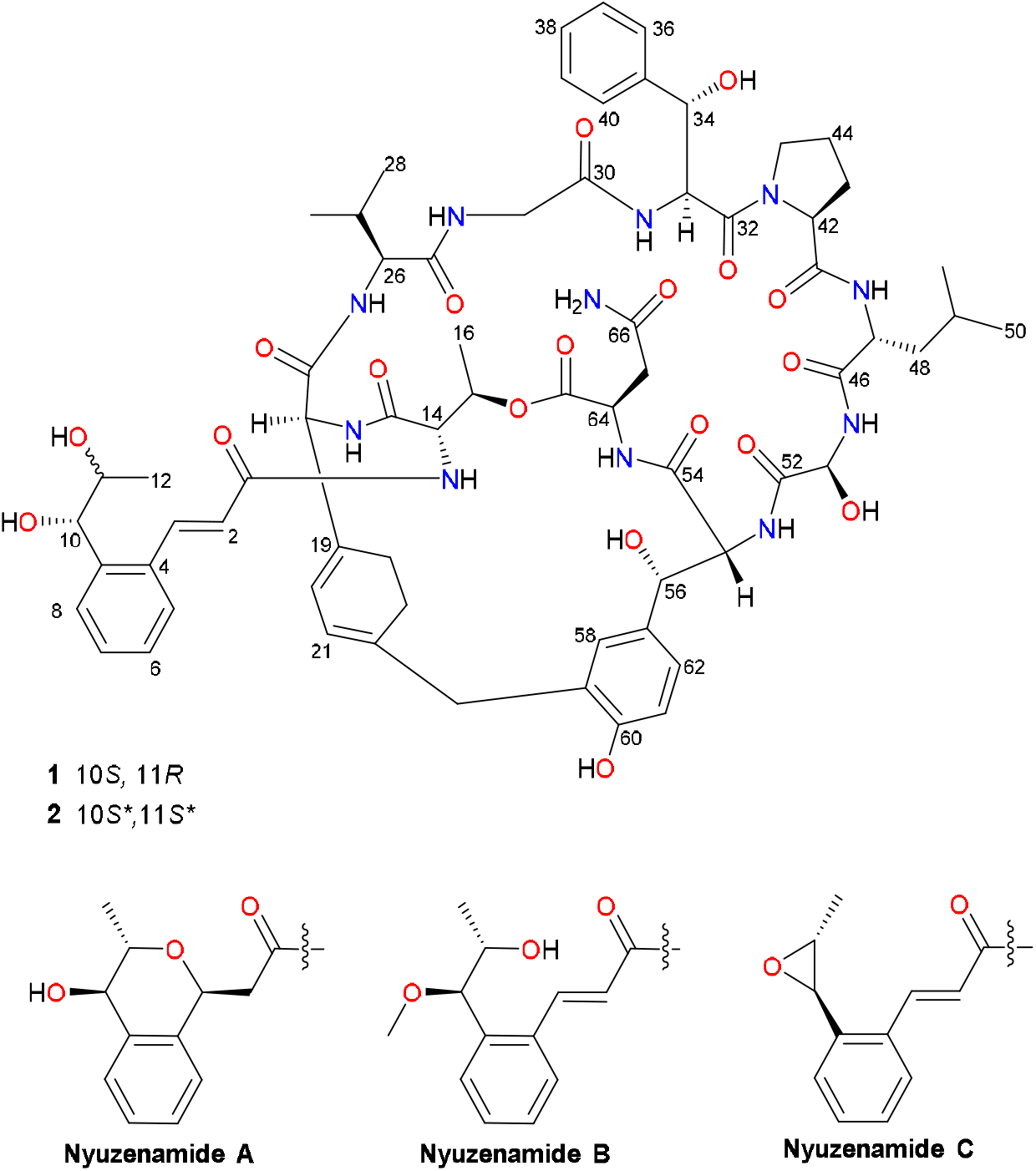
Chemical structures of nyuzenamides D (**1**), E (**2**) and the reported nyuzenamides A–C.

## RESULTS AND DISCUSSION

To obtain enough materials and confirm the chemical structures, *S. hygroscopicus* subsp. *hygroscopicus* DSM 40578 was cultivated on 5 L of MYM agar and sequentially extracted with EtOAc: *i*PrOH (3:1) (1% formic acid) and MeOH. The extract was then subjected to silica gel column chromatography, followed by reversed-phase column chromatography and HPLC, resulting in the isolation of nyuzenamides D (**1**) and E (**2**) (**Chart 1**).

Nyuzenamide D (**1**) was isolated as optically active colorless needles. The molecular formula was predicted to be C_66_H_81_N_11_O_20_ through HRESIMS analysis (*m/z* 1348.5727 [M+H]^+^). The IR spectrum showed characteristic absorption bands at 3290 and 1653 cm^-1^, indicating the presence of NH or OH and carbonyl functionalities, respectively. The results from the ^13^C NMR and the edited HSQC spectra (**Table 1**) revealed a total of 66 carbons, including 12 carbonyl carbons, 26 aromatic or olefinic carbons, 16 nitrogen or oxygen-bound methine carbons, and 12 aliphatic carbons. The ^1^H NMR spectrum (**Figure S1**) exhibited signals corresponding to 17 exchangeable protons. Correlations from the protons to the respective nitrogen atoms as observed in ^1^H-^15^N HSQC and ^1^H-^15^N HMBC spectra (**Figure S7–S8**), indicated that 11 of these exchangeable protons were amide protons. The remaining signals resonating were then deduced to be hydroxyl protons.

**Table 1.**
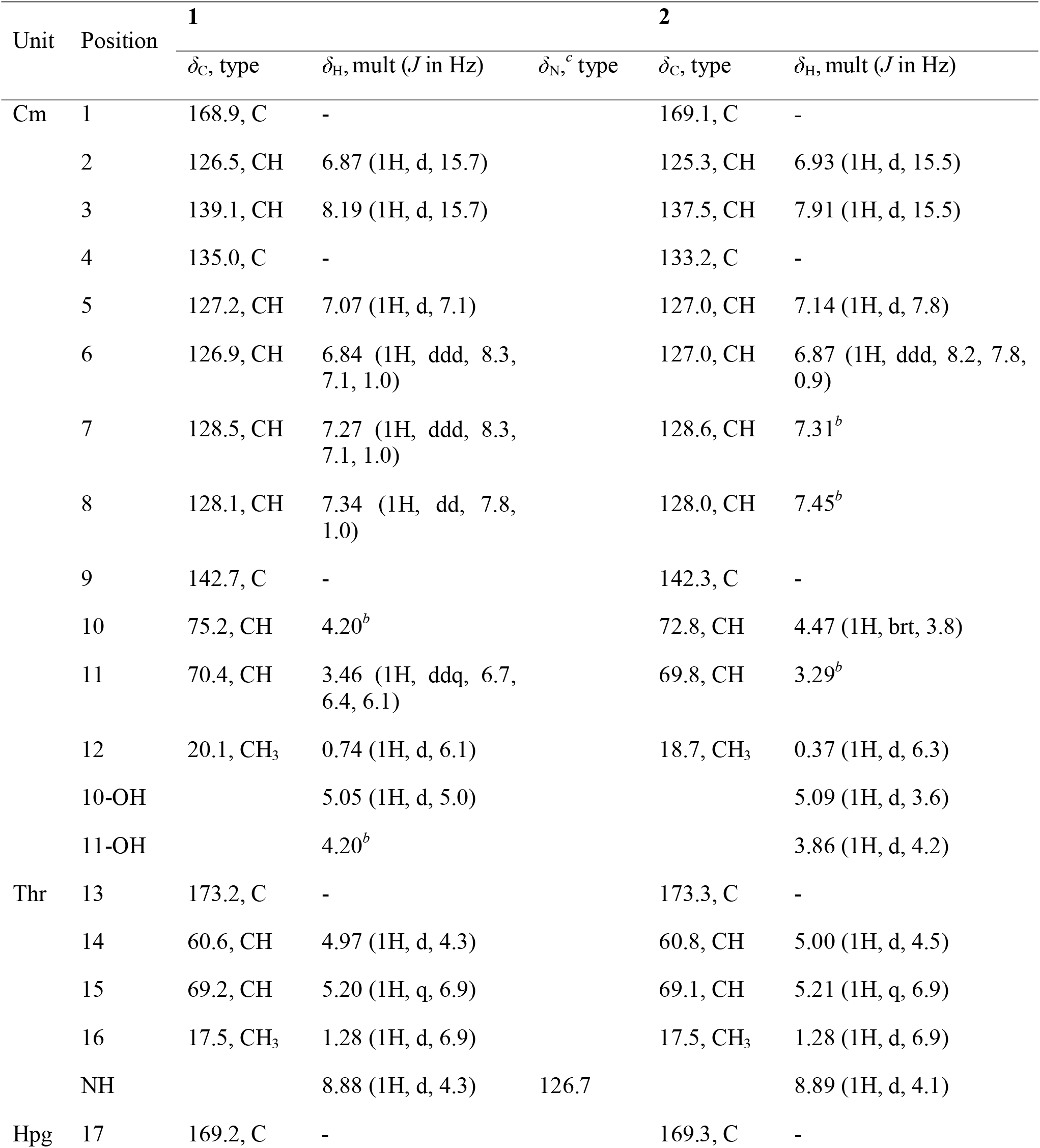
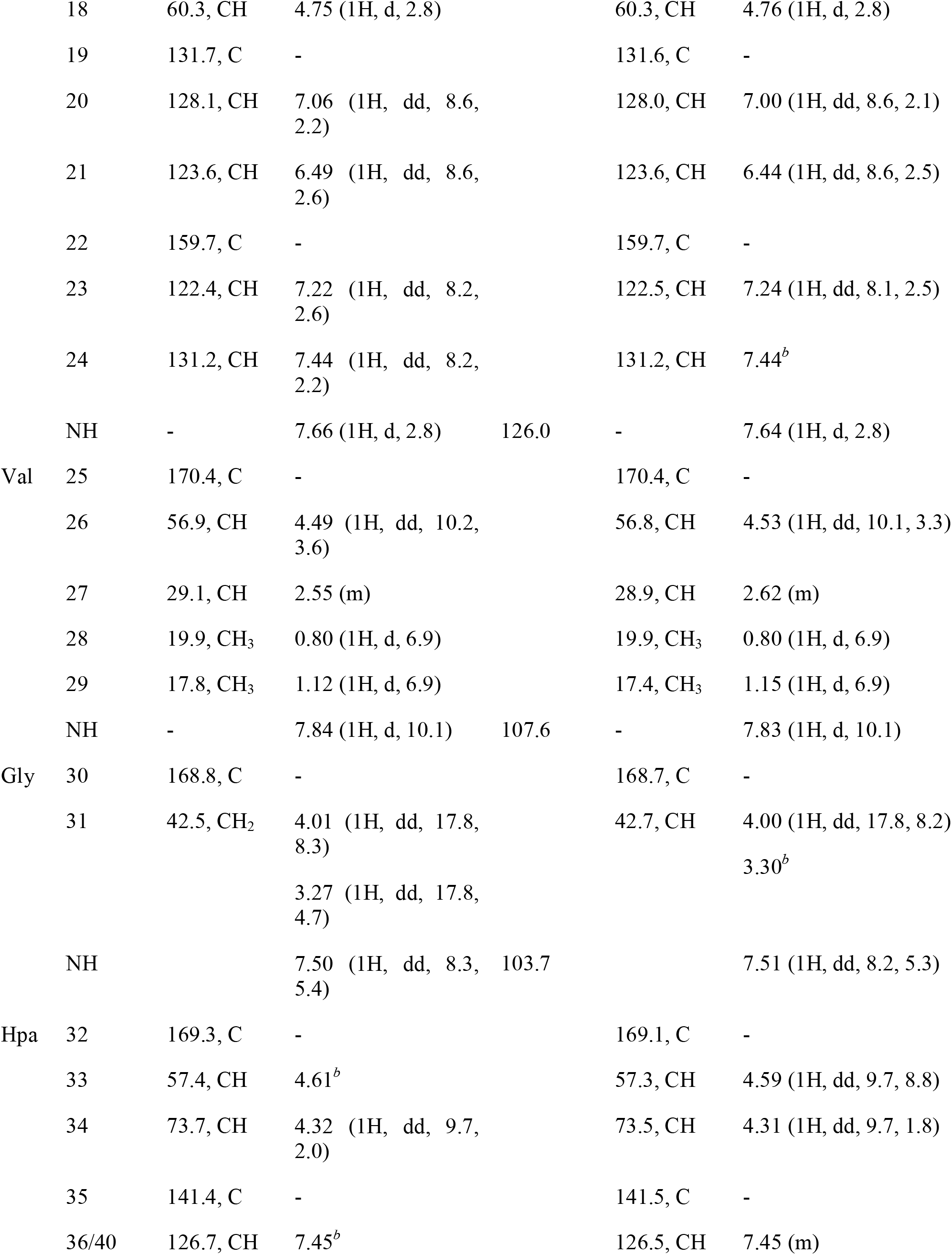
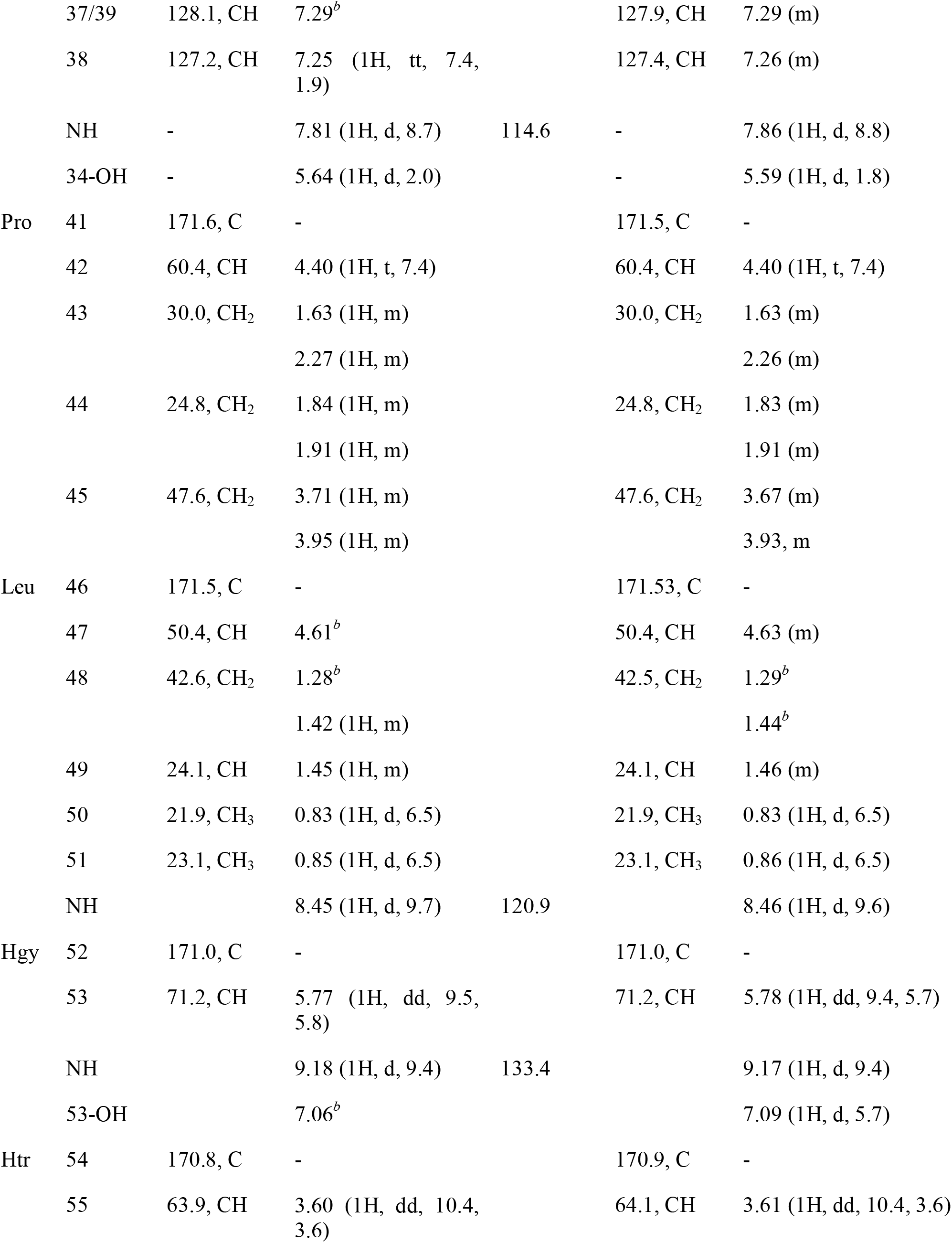
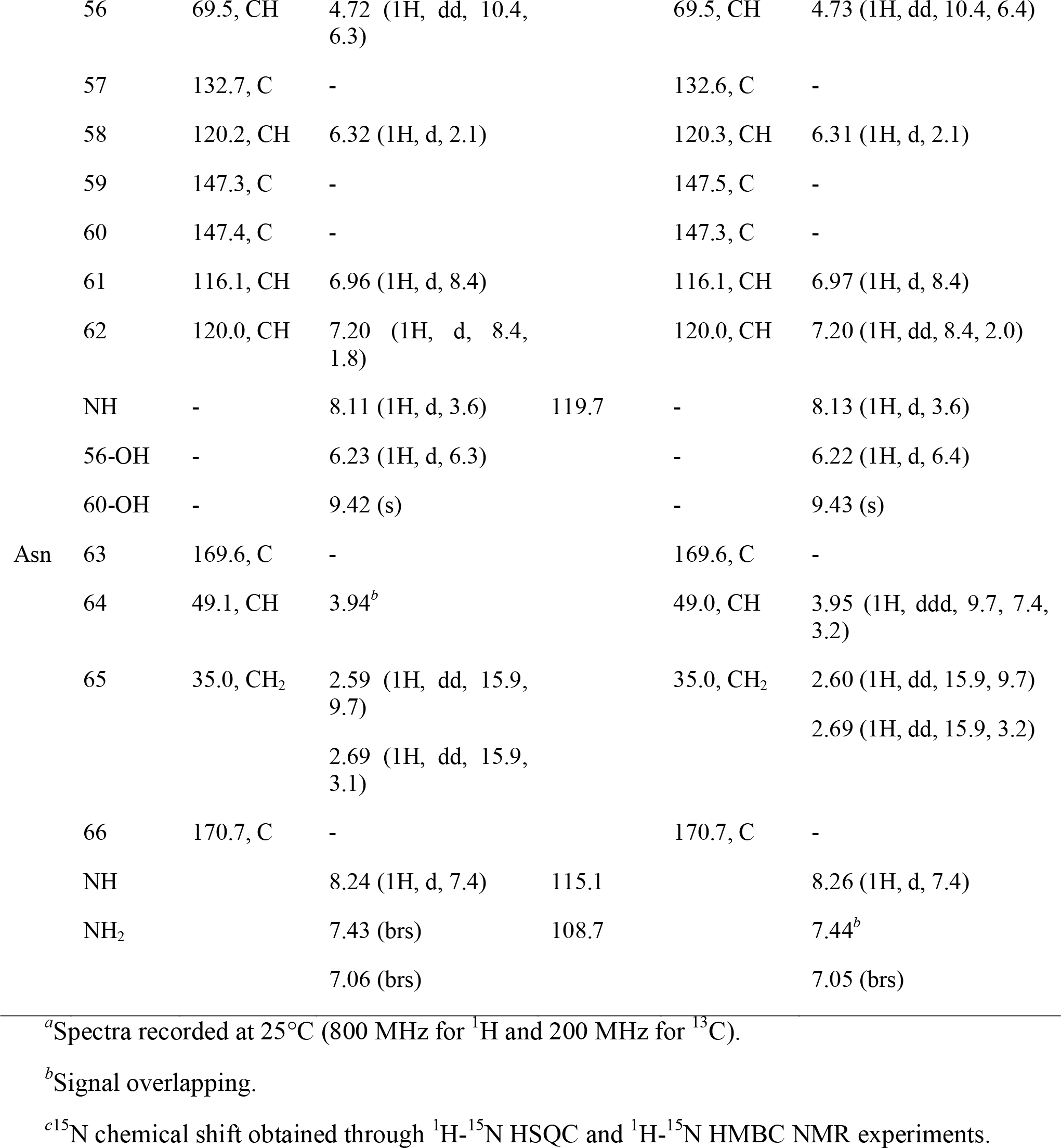
^1^H and ^13^C NMR data for compounds **1** and **2** in DMSO-*d*_6_^a^

Detailed COSY, HMBC and NOESY spectral data analysis (**Figure 1**) allowed the assignment of six proteinogenic amino acids, which include threonine (Thr), valine (Val), glycine (Gly), proline (Pro), leucine (Leu) and asparagine (Asn). The COSY correlations between the amide proton (δ_H_ 9.18), oxymethine (δ_H_ 5.77), and hydroxyl proton (δ_H_ 7.06), along with the HMBC correlations from δ_H_ 5.77 (Hgy-H53) and 7.06 (Hgy-53-OH) to δ_C_ 171.0 (Hgy-C52) suggested the presence of a nonproteinogenic amino acid hydroxyglycine (Hgy). The chemical structure of the *b*-hydroxyphenylalanine (Hpa) was determined through the COSY correlations between δ_H_ 7.81 (Hpa-NH), 4.61 (Hpa-H33), 4.32 (Hpa-H34), and 5.64 (Hpa-34-OH), together with HMBC correlations from δ_H_ 4.32 to the aromatic carbons Hpa-C35 and Hpa-C36/ Hpa-C40, and from the *ortho*-aromatic protons δ_H_ 7.45 to Hpa-C38.

**Figure 1.**
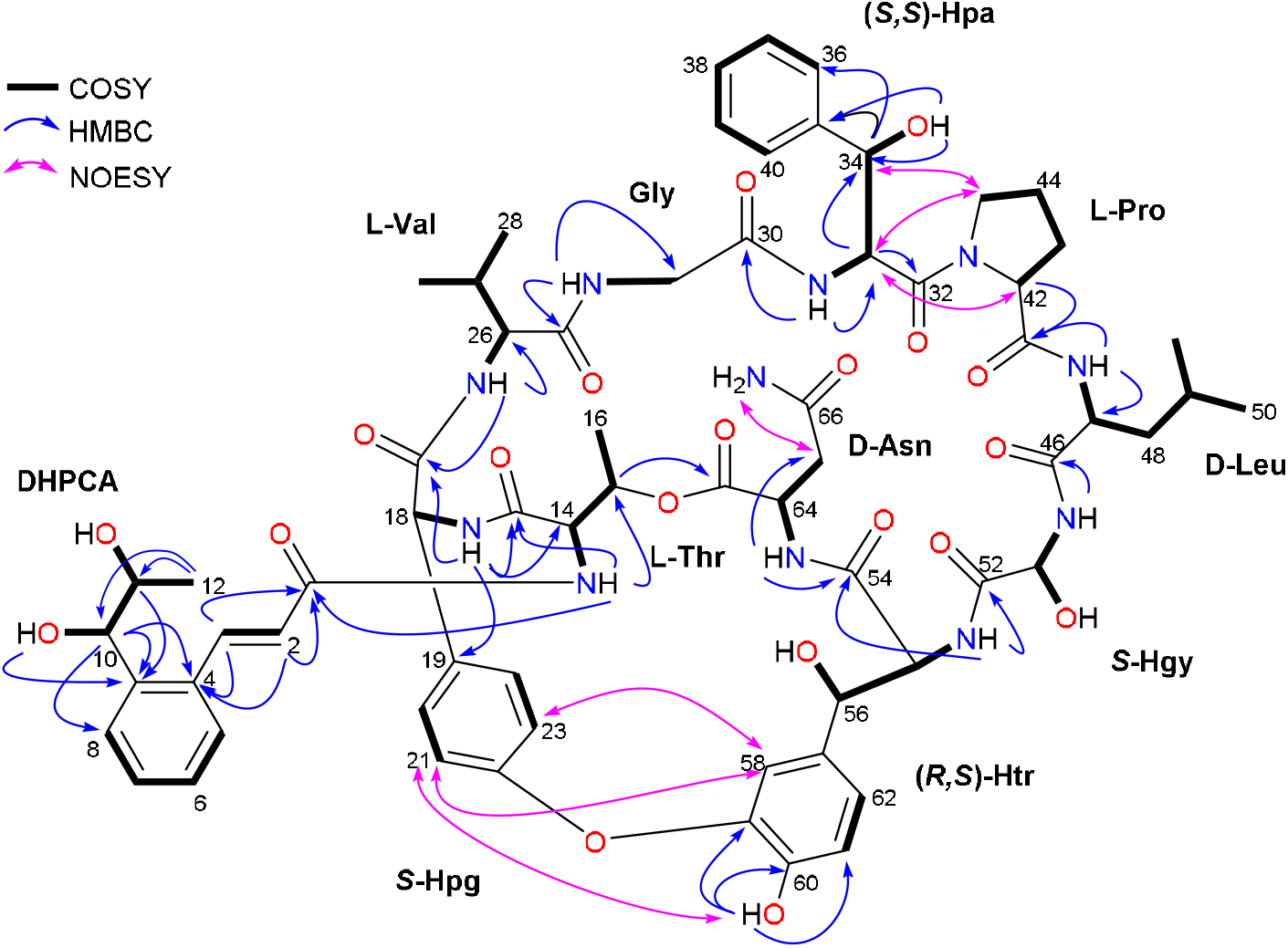
Key COSY, HMBC and NOESY correlations of nyuzenamides D (**1**) and E (**2**).

Based on the COSY spectrum analysis, the consecutive correlations between δ_H_ 8.11 (Htr-NH), 3.60 (Htr-H55), 4.72 (Htr-H56), 6.23 (Htr-56-OH), and the HMBC correlations from the *b*-oxymethine δ_H_ 4.72 to the aromatic carbons δ_C_ 132.7 (Htr-C57), 120.2 (Htr-C58) and 120.0 (Htr-C62), indicated that this amino acid contained an aromatic ring. Further analysis of the HMBC correlations assigned the chemical structure of this amino acid as *b*-hydroxytyrosine (Htr). Similarly, another aromatic ring-containing amino acid residue was determined to be 4-hydroxyphenylglycine (Hpg) following HMBC and COSY data analyses.

After assignment of the amino acid residues, analysis of the remaining protons through COSY spectrum identified three spin systems. The first spin system observed between δ_H_ 6.87 (Cm-H2) and 8.19 (Cm-H3), with a ^3^*J*-coupling constant of 15.7 Hz, supported the presence of a disubstituted *E*-olefin. In the HMBC spectrum, these olefinic protons correlated to a carbonyl carbon (δ_C_ 168.9) and three aromatic carbons at δ_C_ 135.0 (Cm-C4), 127.2 (Cm-C5), and 142.7 (Cm-C9), suggesting a 2-propenoyl unit was conjugated to an aromatic ring. An *ortho*-disubstituted benzene ring was deduced by COSY and HMBC correlations. Furthermore, a 2-propenoyl unit was linked to the aromatic ring at Cm-C4 position, thus forming a cinnamoyl moiety. The last fragment was determined as 1,2-dihydroxypropyl group following COSY correlations, and subsequent analysis on the HMBC spectrum revealed that this vicinal-dihydroxy unit was connected to the cinnamoyl moiety. Hence, the structure of this nonpeptidic moiety was assembled to 2-(1,2-dihydroxypropyl) cinnamic acid (DHPCA).

The HMBC correlations from amide protons to the adjacent amide carbons established the two amino acids sequences: Asn-Htr-Hgy-Leu-Pro and Hpa-Gly-Val-Hpg-Thr-DHPCA. The NOE (NOESY) correlations of Hpa-H33 with Pro-H42, Pro-H43, Pro-H44, and Pro-H45 connected the two sequences via an amide linkage between Pro and Hpa. The HMBC correlation from Thr oxymethine δ_H_ 5.20 to a carboxyl carbon δ_C_ 169.6 (Asn-C63) indicated that Thr and Asn were connected via an ester linkage. Consequently, the planar structure of **1** was assigned as nyuzenamide D, a 2-(1,2-dihydroxypropyl) cinnamic acid containing-bicyclic decadepsipeptide. To definitively assign the absolute configurations, crystallography studies were carried out on compounds **1** and **2**. Slow evaporation of **1** in MeOH at room temperature yielded colorless needles suitable for a single-crystal X-ray diffraction study (**Figure 2**). The analysis of crystal data confirmed the absolute configurations of the proteinogenic amino acids as D-Asn, D-Leu, L-Pro, L-Thr, and L-Val. The unusual amino acids were identified as (*S*)-Hpg, (2*S*, 3*S*)-Hpa, (*S*)-Hgy, (2*R*, 3*S*)-Htr, and the 1,2-dihydroxypropyl cinnamic acid was determined to have (10*S*, 11*R*) configurations.

**Figure 2.**
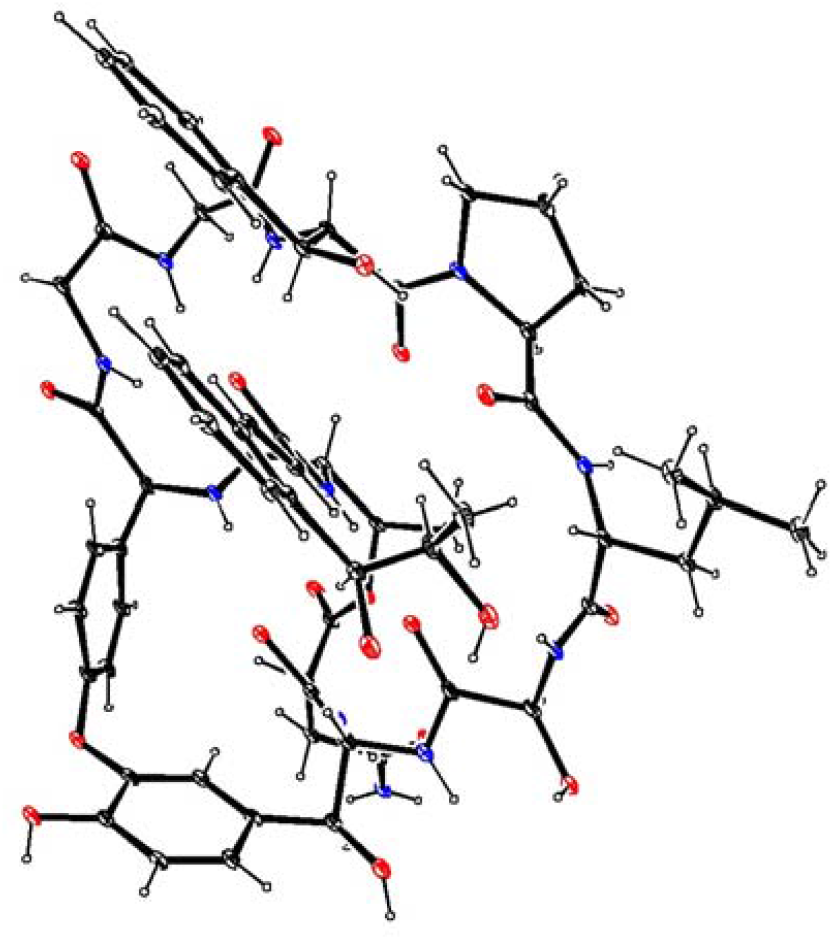
ORTEP drawing of nyuzenamide D (**1**). The methanol molecule has been omitted for clarity.

Nyuzenamide E (**2**) was obtained as a white amorphous powder, and the molecular formula was predicted as C_66_H_81_N_11_O_20_ with 32 index of hydrogen deficiency, based on HRESIMS data with m/z 1348.5726 [M+H]^+^. Systematic analysis of ^1^H, ^13^C, and 2D NMR data revealed that **2** consisted of six proteinogenic amino acids (Thr, Val, Gly, Pro, Leu, Asn) and four unusual amino acids (Hpg, Hpa, Hgy, Htr), which are identical to the peptidic backbone of compound **1**. The amino acid sequence was confirmed through a comprehensive set of HMBC and NOESY correlations. Notably, the ^1^H and ^13^C chemical shifts of the DHPCA unit in **2** differed from those in **1** (**Table 1**). For instance, the resonances of Cm-C10 (δ_c_ 72.8) and Cm-H12 (δ_H_ 0.37) in **2** were more shielded compared to **1** (δ_C_ 75.2 and δ_H_ 0.74). Hence, it is suggested that the deviation may arise from different configurations in the two stereocenters, Cm-C10 and Cm-C11, within the 1,2-dihydroxylpropyl chain. Similar observation was reported previously, where different ^1^H and ^13^C chemical shifts were noted in the (*S,S*)- and (*R,R*)-1,2-epoxypropyl cinnamic acid (EPCA) moieties of epoxinnamide and nyuzenamide C, respectively.^10,16^

In comparison to the coupling constants and NOESY correlations, and from a biosynthetic perspective, it can be concluded that the absolute configurations of the common and nonproteinogenic amino acids in **2** were identical to those in **1**. However, the difference in the chemical shifts at the 1,2-dihydroxypropyl group of both compounds indicated that **2** is a diastereomer of **1**. According to the predicted biosynthetic pathway for a pre-NRPS assembled vicinal-dihydroxy cinnamic acyl chain in a cyclodepsipeptide, atrovimycin,^5^ the formation of this moiety involves multiple enzyme-catalyzed steps, including the hydroxylation of the intermediate EPCA moiety by epoxide hydrolase to vicinal diol. Epoxide hydrolases function through the nucleophilic attack of OH^-^ on the carbon atom, leading to the opening of the epoxide. The absolute configuration of the resulting diol can be retained or inverted, depending on the substitutional pattern of carbon atom involved and the regioselectivity of the enzyme.^17^ Therefore, based on the crystal and NMR spectroscopic data, the chiral centers of the vicinal dihydroxy group in the cinnamic acid moiety of **2** were assigned as 10*S*\*, *11S*\*.

Due to the incomplete genome data of *S. hygroscopicus* subsp. *hygroscopicus* DSM 40578, a comprehensive whole-genome resequencing was conducted using the Oxford Nanopore Technologies MinION and Illumina NovaSeq systems. The biosynthetic gene cluster (BGC) annotation was carried out by antiSMASH 7.0^18^ and a total of 42 putative BGCs were detected (**Table S19**). Notably, the putative BGC number 22 exhibits high similarity to previously speculated biosynthetic gene clusters thought to be responsible for nyuzenamide C biosynthesis in *Streptomyces* sp. DM14.^10^ AntiSMASH analysis of this core BGC revealed three genes (*nyuO, nyuP* and *nyuS*) coding for ten modular NRPSs (**Figure 3**), which are proposed to be responsible for incorporating ten different amino acid residues. Analysis of the A domains indicated the incorporation of Thr (A1), X (A2), Val (A3), Gly (A4), Phe (A5), Pro (A6), Leu (A7), Gly (A8), Tyr (A9) and Asn (A10). Two epimerase domains in modules 7 and 9 fit well with the presence of one D-Leu and D-Tyr residues in the assigned chemical structures, respectively, whereas glycine in module 4 does not have an absolute configuration. Biosynthesis of cinnamoyl moiety has been reported in atratumycin^4^ and skyllamycin.^19^ Nine gene analogues (**Figure 3**) to the cinnamate biosynthetic gene cassettes were also observed in the *nyu* BGC. Moreover, three P450 oxygenases are proposed to catalyze reactions including the oxygenation bridging Hpg and Htr, *b* hydroxylation of phenylalanine and tyrosine, and *a* hydroxylation of glycine.

**Figure 3.**
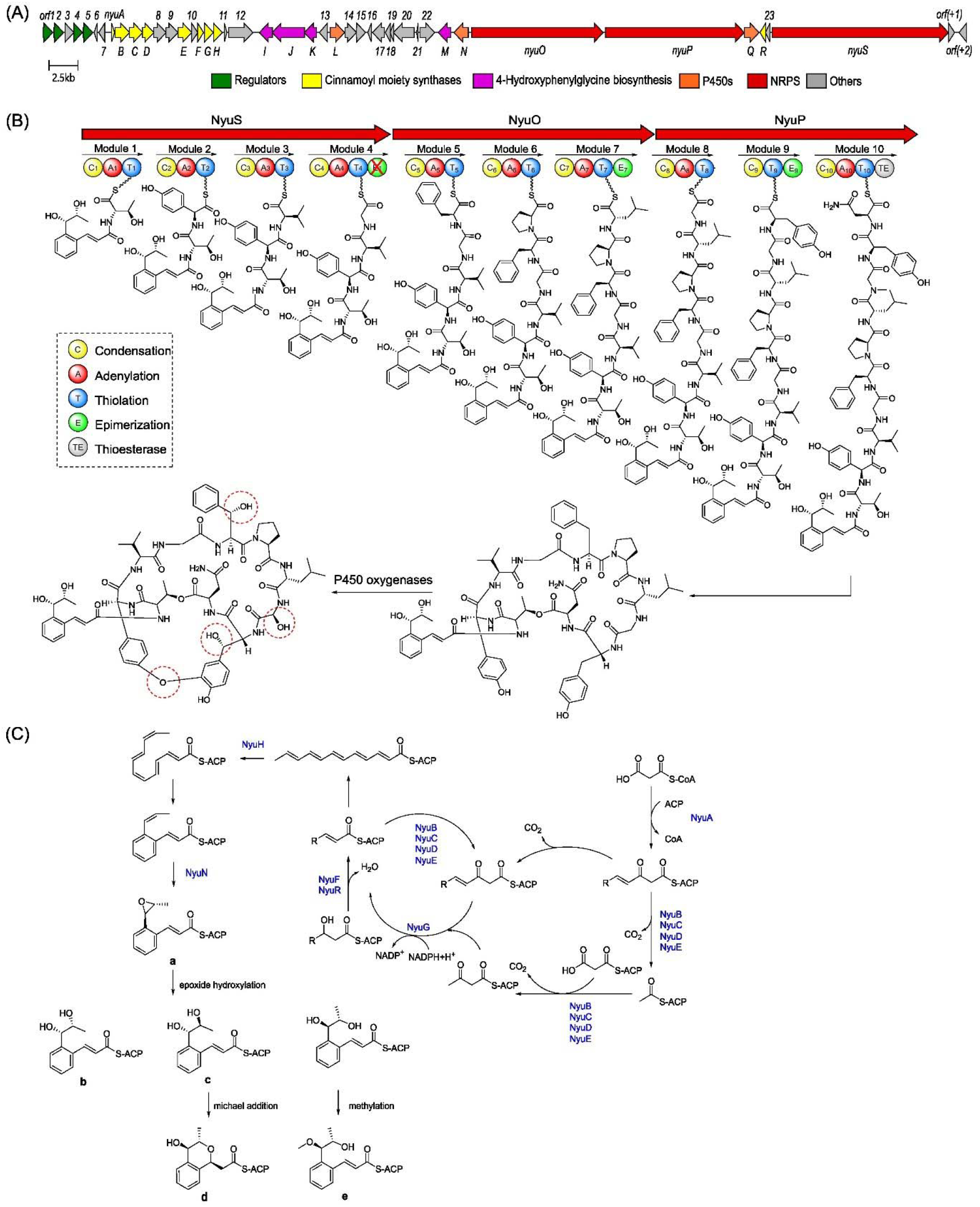
Biosynthetic gene cluster of nyuzenamides D (**1**) and (**2**). (A) Gene organization of *nyu* BGC. (B) Schematic representation of biosynthesis of **1** by *S. hygroscopicus* subsp. *hygroscopicus* DSM 40578. C, condensation domain; A, adenylation domain; T, peptide-carrier protein; E, epimerase; TE, thioesterase. (C) Proposed biosynthetic pathway for the 2-(1,2-dihydroxypropyl) cinnamic acid acyl chain, cinnamoyl units a–e corresponding to the precursors of nyuzenamides A–E.

To experimentally verify the role of the putative nyuzenamides BGC *in vivo*, the CRISPR-Cas9 genome editing system^20^ was introduced into *S. hygroscopicus* DSM 40578 to inactivate the expression of the core polyketide synthetase NyuS (DmlA in the nyuzenamide C BGC from *Streptomyces* sp. DM14).^10^ The HRESIMS analysis showed the production of nyuzenamides D and E was abolished in the D*nyuS* mutant, as shown in **Figure 4**. Through the integration of both in silico and in vivo data, we have conclusively established that this BGC is pivotal for the biosynthesis of nyuzenamides D (**1**) and E (**2**) in *S. hygroscopicus*.

**Figure 5.**
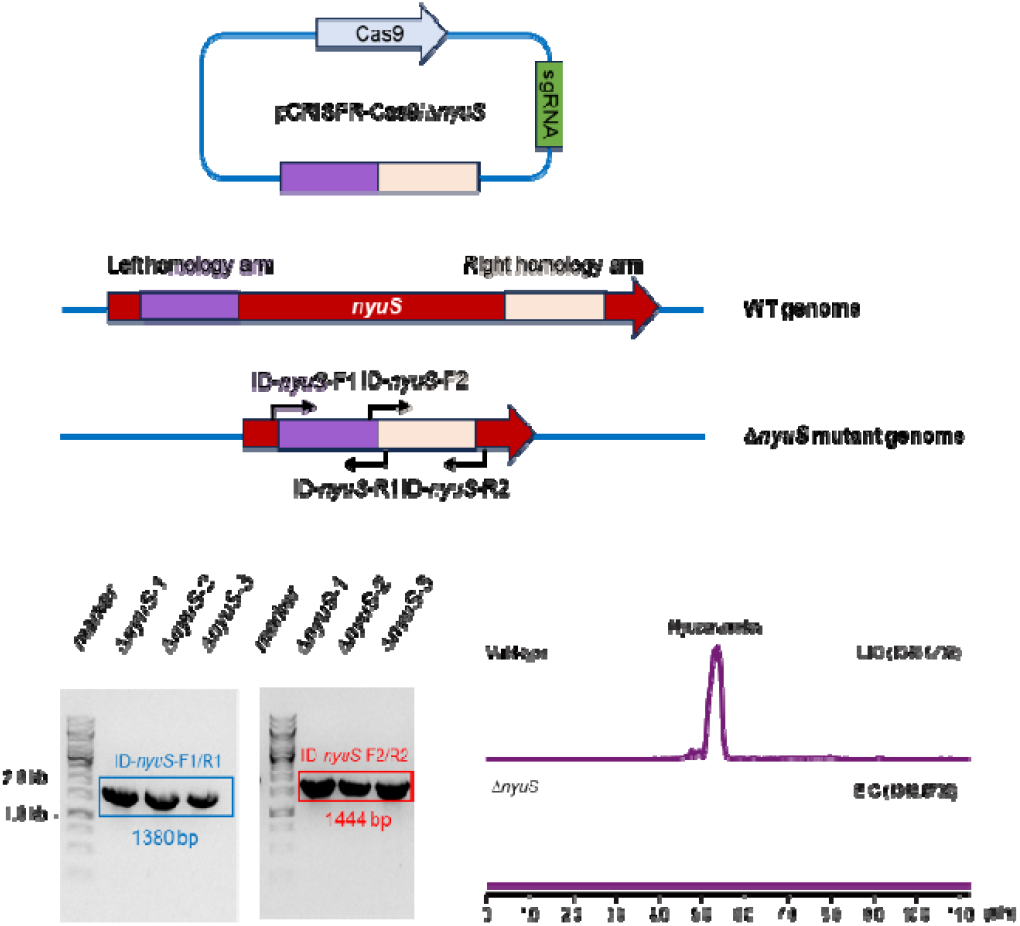
CRISPR-based gene deletion of *nyuS* in *S. hygroscopicus* subsp. *hygroscopicus* DSM 40578. Extracted ion chromatogram (EIC) for **1** and **2** (*m/z* 1348.5732 [M+H]^+^) in the wild type strain and the mutant *S. hygroscopicus*/Δ*nyuS*.

Nyuzenamides A–E share the same cyclic nonribosomal peptide core structure. Nyuenamides A–C were reported to exhibit cytotoxicity or antifungal activities.^8-9^ In the present study, nyuzenamides D (**1**) and E (**2**) were evaluated for their antibacterial and antifungal activities against *Bacillus subtilis, Staphylococcus aureus, Pseudomonas aeruginosa* PAO1, *Candida albicans* IBT 654, *Alternaria solani, Botrytis cinerea, Aspergillus flavus, Aspergillus niger, Aspergillus fumigatus*, and *Aspergillus tubingensis*; none of the compounds showed inhibitory activity at tested concentration (MIC > 64 μg/mL). Additionally, neither compound **1** nor **2** showed significant cytotoxic activity against the human prostate cancer cell lines LNCaP and C4-2B (IC_50_ >10 μM) under the tested conditions (72-hour continuous exposure to various concentrations, up to 10 μM). Apparently, changes in the cinnamoyl moiety, especially substitutions at C-10 and C-11, led to the differences of bioactivity among nyuzenamides.

## EXPERIMENTAL SECTION

### General Experimental Procedures

Specific rotations were acquired using an AUTOPOL III Automatic polarimeter. IR data were acquired on Bruker Alpha FTIR spectrometer using OPUS version 7.2. NMR spectra were acquired on 800 MHz Bruker Avance III spectrometer equipped with a TCI CryoProbe using standard pulse sequences. The ^1^H and ^13^C NMR chemical shifts were referenced to the residual solvent peaks at δ_H_ 2.50 and δ_C_ 39.5 ppm for DMSO-*d*_6_. NMR data were processed using MestReNova 14.1.1. UHPLC-HRESIMS was performed on an Agilent Infinity 1290 UHPLC system equipped with a diode array detector. UV-vis spectra were recorded from 190 to 640 nm. Silica gel (40–63 μm, 60 Å, Sigma-Aldrich) was packed into a 25 g Biotage SNAP cartridge and Phenomenex Luna C18(2) 100 Å (11–14 μm) silica was packed into a 50 g cartridge for normal and reversed-phase flash column chromatography, respectively. Biotage Isolera One Flash Chromatography system was used for flash chromatography. All solvents and chemicals used for UHPLC-HRESIMS were VWR Chemicals LC-MS grade, while for metabolites extraction and chromatography, the solvents were of VWR Chemicals HPLC grade. H_2_O was filtered using a Mili-Q ultrapure water system.

### Bacterial Cultivation

*S. hygroscopicus* subsp. *hygroscopicus* DSM 40578 was obtained from German Collection of Microorganisms and Cell Cultures GmbH. The spore suspension of *S. hygroscopicus* was inoculated on ISP2 agar (4 g/L yeast extract, 10 g/L of malt extract, 4 g/L dextrose, 20 g/L agar, and 1 L of distilled water, pH 7.2) for 3 days. The seed culture was prepared by transferring the agar plugs to a 200 mL conical flask containing 50 mL of ISP2 liquid media and incubated for 5 days at 28 °C with shaking at 180 rpm. 10 mL of the seed culture was then inoculated on 5 × 1 L of MYM agar trays (4 g/L maltose, 4 g/L yeast extract, 10 g/L malt extract, 10 g/L agar, 0.5 L of tap water and 0.5 L of distilled water, pH 7.0–7.4) and incubated at 30 °C for 7 days.

### UHPLC-HRESIMS sample preparation and analysis

An agar plug (6 mm diameter) of the bacterial culture was transferred to a vial (Eppendorf) and extracted with 1 mL of isopropanol: ethyl acetate 1:3 (v/v) with 1% formic acid under ultrasonication for 15 minutes. The extracts were then transferred to new Eppendorf vials, evaporated to dryness under N_2_, and re-dissolved in methanol to meet the final concentration of 10 mg/mL. After centrifugation at 10000 rpm for 3 min, the supernatants were transferred to HPLC vials, diluted to 1 mg/mL with methanol and subjected to ultrahigh-performance liquid chromatography-high resolution electrospray ionization mass spectrometry (UHPLC-HRESIMS) analysis. Other samples during isolation and purification process were all prepared with methanal and diluted to 1 mg/mL for UHPLC-HRESIMS analysis. HR-ESI-MS data were acquired on an Agilent Infinity 1290 UHPLC system equipped with a diode array detector and coupled to an Agilent 6545 QTOF MS equipped with Agilent Dual Jet Stream ESI. Separation was achieved on a 250 × 2.1 mm i.d., 2.7 μm, Poroshell 120 Phenyl Hexyl column (Agilent Technologies) held at 40 °C. The sample, 1 μL, was eluted at a flow rate of 0.35 mL min^−1^ using a linear gradient from 10% acetonitrile/water buffered with 20 mM formic acid to 100% acetonitrile in 15 min, held for 2 min and equilibrated back to 10% acetonitrile/water in 0.1 min. Starting conditions were held for 3 min before the following run. The MS settings were as follows: drying gas temperature of 160 °C, a gas flow of 13 L min^−1^, the sheath gas temperature of 300 °C and flow of 16 L min^−1^. Capillary voltage was set to 4000 V and nozzle voltage to 500 V in positive mode. All data were processed using Agilent MassHunter Qualitative Analysis software (Agilent Technologies, USA). All solvents used for chromatography and HR-MS were purchased from VWR Chemicals with LC-MS grade, while for metabolites extraction, the solvents were of HPLC grade.

### Extraction and Isolation

The cultured agar was extracted twice with EtOAc: *i*PrOH (3:1) (1% formic acid) and methanol. The dried acidic EtOAc: *i*PrOH extract (7 g) was divided into two portions, pre-adsorbed onto Si-gel, and chromatographed over a Si-gel flash column. The column was flushed with a 2% stepwise gradient solvent system from 100% CH_2_Cl_2_ to 90% CH_2_Cl_2_/10% MeOH, followed by 80% CH_2_Cl_2_/20% MeOH and further eluted with a 10% stepwise gradient from 50% CH_2_Cl_2_/50% MeOH to 100% MeOH at a flow rate of 25 mL/min, which resulted in 24 fractions. Based on silica TLC and UV profiles (254 and 280 nm), similar fractions were combined to afford 10 fractions (F1–F10). Fraction F4 (2.34 g) was pre-adsorbed onto Phenomenex C18(2) silica and further fractionated by RP C_18_ flash column chromatography. A linear gradient solvent system from 30% MeOH/70% H_2_O (0.1% formic acid) to 100% MeOH (0.1% formic acid) was employed at a flow rate of 50 mL/min, and 14 fractions (F1–F14) were collected. Based on (+)-HRESIMS analysis, fraction F12 (90.9 mg) was purified using a semipreparative phenyl-hexyl HPLC column (5 μm, 100 Å, 250 × 10 mm). A linear gradient from 50% MeOH/50% H_2_O (0.01 % TFA) to 90% MeOH/10% H_2_O (0.01 % TFA) at a flowrate of 4 mL/min was run over 40 min to yield the semipure nyuzenamide D (**1**, 8.5 mg, *t*_R_ 19–20 min) and nyuzenamide E (**2**, 4.9 mg, *t*_R_ 20–21min). Further purification of **1** by a semipreparative phenyl-hexyl HPLC column with a linear gradient from 35% MeOH/65% H_2_O (0.01 % TFA) to 80% MeOH/20% H_2_O (0.01 % TFA), at a flowrate of 4 mL/min for 20 min afforded nyuzenamide D (**1**, 4.8 mg, *t*_R_ 5–6 min).

***Nyuzenamide D (1)*:** colorless needles;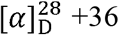; UV (MeCN/H_2_O) λ_max_ 216, 278 nm; IR (ATR) v_max_ 3290, 1653, 1541, 1526 cm^-1^; ^1^H and ^13^C NMR data, see Table 1; (+)-HRESIMS *m*/*z* 1348.5727 [M+H]^+^ (Calcd for C_66_N_11_O_20_H_82_, 1348.5732).

***Nyuzenamide E (2)*:** white, amorphous powder;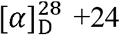; UV (MeCN/H_2_O) λ_max_ 218, 280 nm; IR (ATR) v_max_ 3290, 1655, 1526, 1439 cm^-1^; ^1^H and ^13^C NMR data, see Table 1; (+)-HRESIMS *m*/*z* 1348.5726 [M+H]^+^ (Calcd for C_66_N_11_O_20_H_82_, 1348.5732).

### X-ray Crystallography Analysis

Single crystals of nyuzenamide D (**1**) were mounted in nylon loops and transported to the PETRA III P14 beamline for data collection. The crystal was kept at 100 K during data collection. Using Olex2,^21^ the structure was solved with the SHELXT^22^ structure solution program utilizing Intrinsic Phasing and refined with the SHELXL^23^ refinement package using Least Squares minimization. Thermal ellipsoid plots were generated using the program ORTEP-3^24^ integrated within the WINGX suite of programs^25^. Thermal ellipsoids are shown at the 5 % level. The crystal data of **1** has been deposited with the Cambridge Crystallographic Data Centre database (https://www.ccdc.cam.ac.uk/) with accession number CCDC No 2349669 and can be obtained free of charge.

### Crystal Data for Nyuzenamide D (1)

C_66_H_81_N_11_O_20_, *M* = 1365.93, monoclinic, a = 25.7170(7) Å, b = 16.4720(7) Å, c = 21.5040(5) Å, α = 90.0°, β = 116.75°, γ = 90.0°, V = 8134.3(5) Å^3^, T = 293(2) K, space group C2, Z = 4, μ (Cu Kα) = 0.079 mm^-1^, 46710 reflections collected, 13203 independent reflections (R_int_ = 0.0306, R_sigma_ = 0.0265). The final R_1_ values were 0.0544 (I > 2σ(I)). The final wR_2_ values were 0.1589 (I > 2σ(I)). The final R_1_ values were 0.0557 (all data). The final wR_2_ values were 0.1608 (all data). The goodness of fit on F^2^ was 1.088. The Flack parameter is 0.37(17).

### Genome Sequencing

The *S. hygroscopicus* subsp. *hygroscopicus* DSM 40578 was growing in 50 mL sterilized liquid ISP2 medium (yeast extract 4.0 g; malt extract 10.0 g; and dextrose 4.0 g in 1.0 L distilled water, pH = 7.2) in 250 mL flask at 30 °C and 160 rpm for 3 days. The genomic DNA was isolated and purified using the QIAGEN Genomic-tip G100 kit according to manufacturer’s instructions. The procedure of genomic DNA sequencing was combining Oxford Nanopore MinION and Illumina MiSeq systems. The de novo assembly was carried out using Flye (v2.9.1 release) and Unicycler (v0.4.8) polishing module. The genome sequence of *S. hygroscopicus* subsp. *hygroscopicus* DSM 40578 was deposited in the National Center for Biotechnology Information (NCBI) under GenBank accession number AB184428.

### Genome Editing

Based on efficient CRISPR-based gene knock-out method described in the previous study,^20^ the modified gene knock-out plasmid pCRISPR-D*nyuS* was constructed and introduced into wild-type *S. hygroscopicus* subsp. *hygroscopicus* DSM 40578 strain by conjugation with donor strain ET12567/pUZ8002 on ISP2 solid medium supplemented with 20 mM MgCl_2_. The exconjugants after resistance screening (50 g mL^-1^ apramycin and 25 g mL^-1^ nalidixic acid) were further verified by PCR reaction.

### Antimicrobial Assays

The antibacterial assay was performed in cationically-adjusted Mueller-Hinton broth (CAMHB), whereas Roswell Park Memorial Institute (RPMI) 1640 medium was used for anticandidal assay according to previously described methods.^26,27^ The minimum inhibitory concentration (MIC) was defined as the lowest concentration that inhibited 90% of bacteria or fungal growth. Antifungal activity against filamentous fungi was evaluated using agar well diffusion assay. Briefly, 64 μg of the compounds (dissolved in 50% CH_2_Cl_2_/50% MeOH) were transferred to their respective well on the potato dextrose agar plate, and the plates were left in the laminar flow for 30 mins to allow the solvent to evaporate. A 5 mm diameter mycelial plug or spore suspension of each fungus was then inoculated on the middle of the plate, and the fungal growth was recorded after incubation at 28 °C for 10 days. Gentamicin and chloramphenicol were used as positive controls for *Pseudomonas aeruginosa, Bacillus subtilis*, and *Staphylococcus aureus*. Amphotericin b was included as a positive control for *Candida albicans, Alternaria solani, Botrytis cinerea, Aspergillus flavus, Aspergillus niger, Aspergillus fumigatus*, and *Aspergillus tubingensis*.

### Cytotoxicity Assay

Compounds **1** and **2** were tested on two different prostate cancer cell lines, LNCaP (RRID:CVCL_0395) and the bone metastatic derivative C4-2B (RRID:CVCL_4784), with cell viability assays performed as previously described by Mohamed et al.^28^

## ASSOCIATED CONTENT

### Data Availability Statement

The NMR data for nyuzenamides D and E have been deposited in the Natural Products Magnetic Resonance Database (NP-MRD; www.np-mrd.org) and can be found at NP0333409 (nyuzenamide D) and NP0333410 (nyuzenamide E). Genomic information is available in NCBI under GenBank GCA_037126205.1 (*Streptomyces hygroscopicus* subsp. *hygroscopicus* DSM 40578).

### Supporting Information

Supporting Information is available free of charge on the ACS Publications website at DOI:

## AUTHOR INFORMATION

Authors

### Author Contributions

### Notes

The authors declare no competing financial interest.

## ACKNOWLEDGMENTS

L.D. acknowledges Novo Nordisk Foundation (BII, PoC NNF20OC0062267, NNF23OC0082881), Carlsberg Infrastructure (CF20-0177) and Innovation Fund Denmark (Grant 1046-00007). K.Y.L thanks the Novo Nordisk Foundation (NNF22OC0079187). The NMR Center DTU and the Villum Foundation are acknowledged for access to the 800 MHz spectrometer. We thank the support from Dr. Tue Sparholt Jørgensen, Dr. Charlotte Held Gotfredsen (DTU NMR Centre) and Dr. Aaron John Christian Andersen (DTU Metabolomics Core).

