## supplemental data for "Discovery and Biosynthesis of Nyuzenamides D and E by Genome Mining in *Streptomyces hygroscopicus*"

**Content**

S1 ^1^H NMR (800 MHz) Spectrum of Nyuzenamide D (**1**) in DMSO-*d_6_*

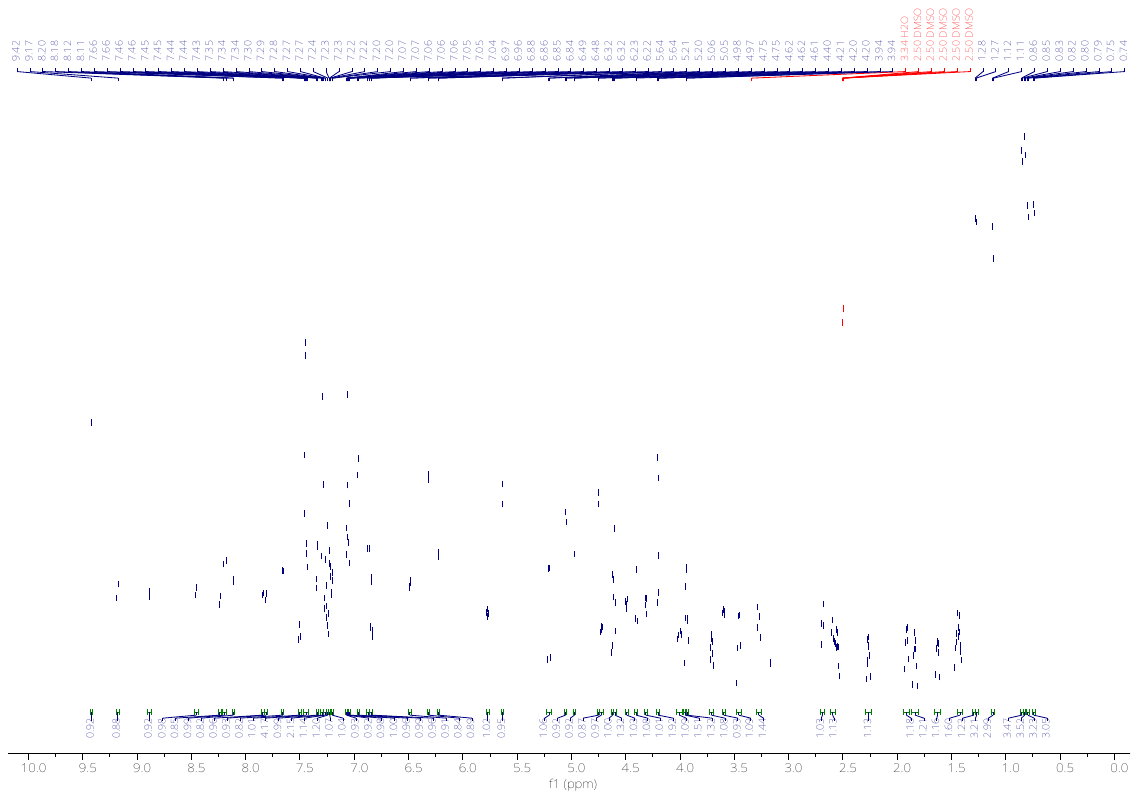

S2 ^13^C NMR (200 MHz) Spectrum of Nyuzenamide D (**1**) in DMSO-*d_6_*

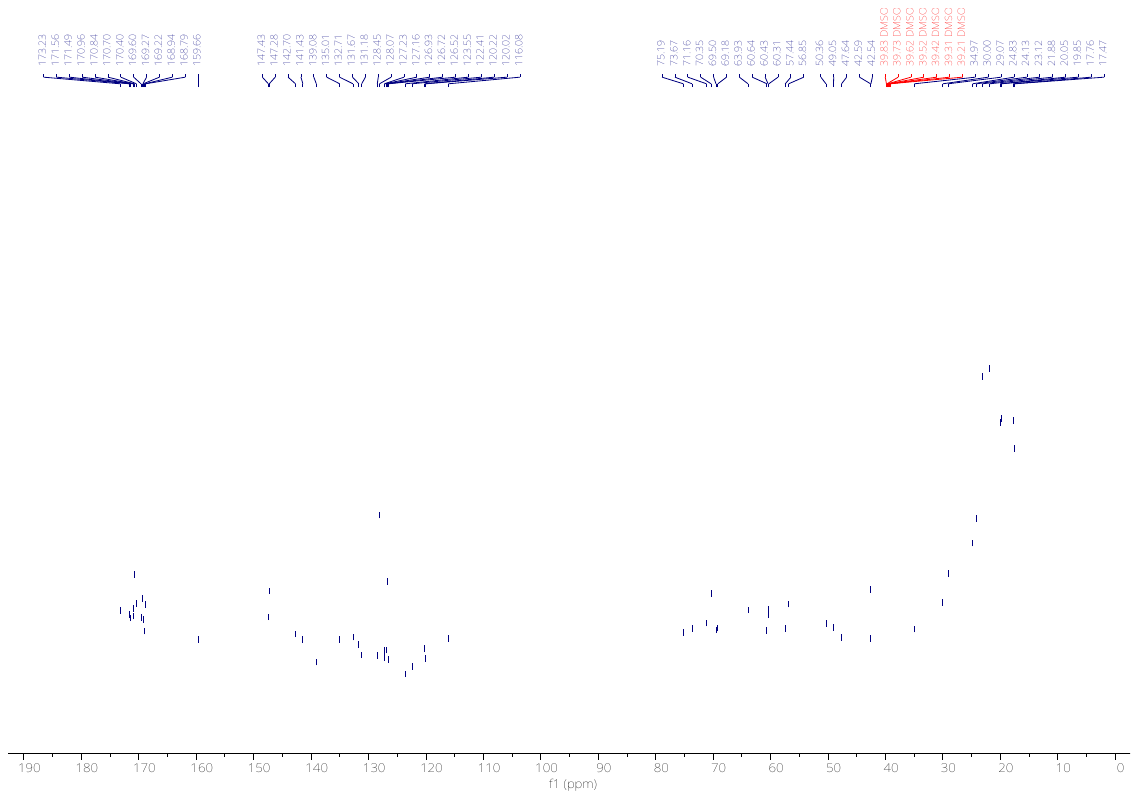

S3 DQF-COSY Spectrum of Nyuzenamide D (**1**) in DMSO-*d_6_*

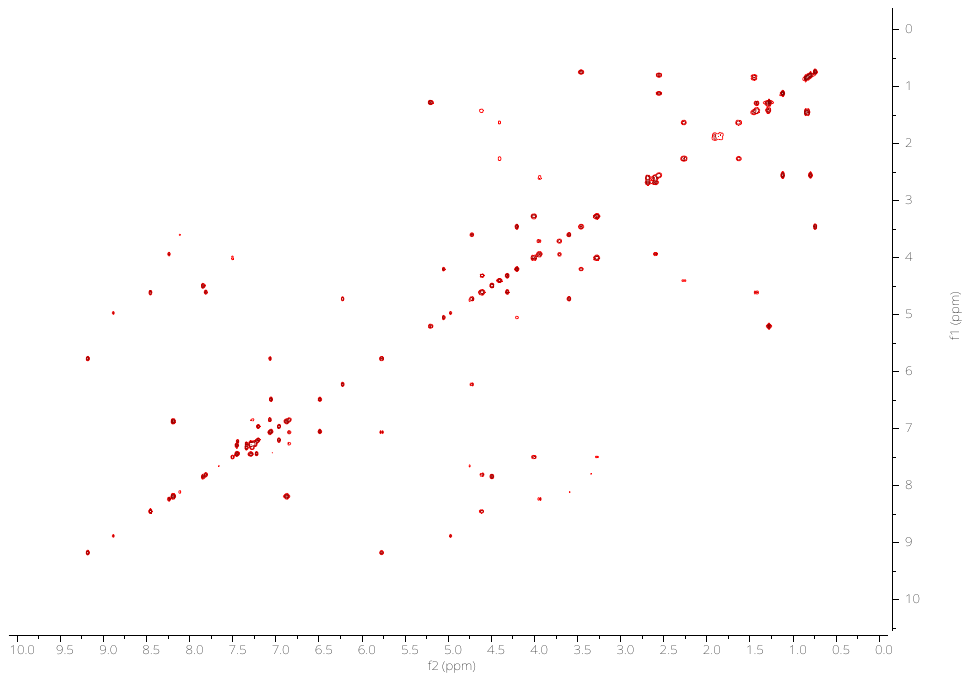

S4 Edited HSQC Spectrum of Nyuzenamide D (**1**) in DMSO-*d_6_*

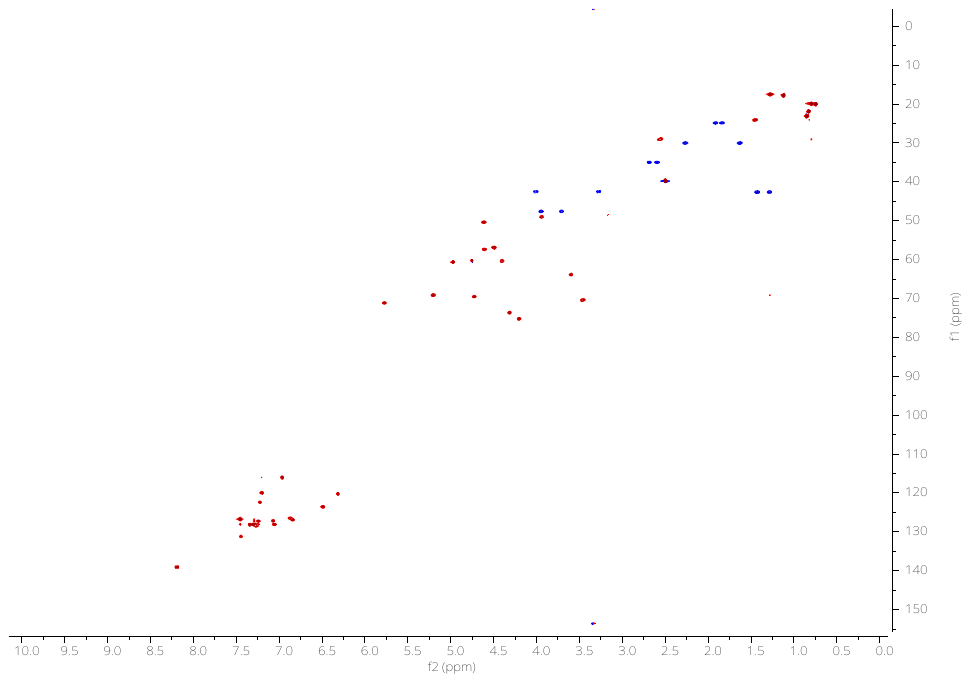

S5 HMBC Spectrum of Nyuzenamide D (**1**) in DMSO-*d_6_*

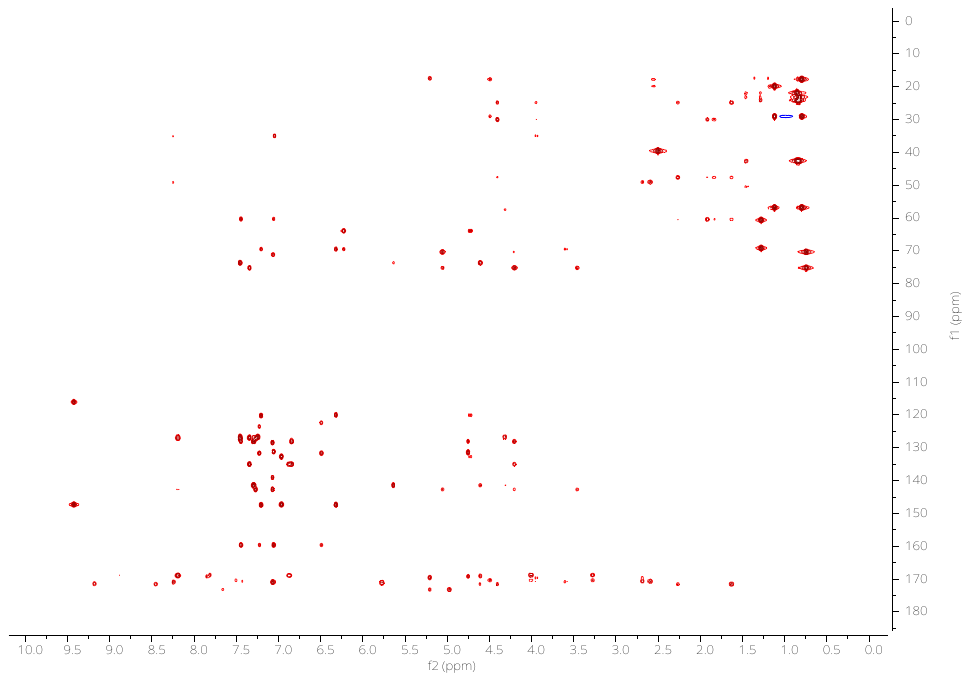

S6 NOESY (600 ms) Spectrum of Nyuzenamide D (**1**) in DMSO-*d_6_*

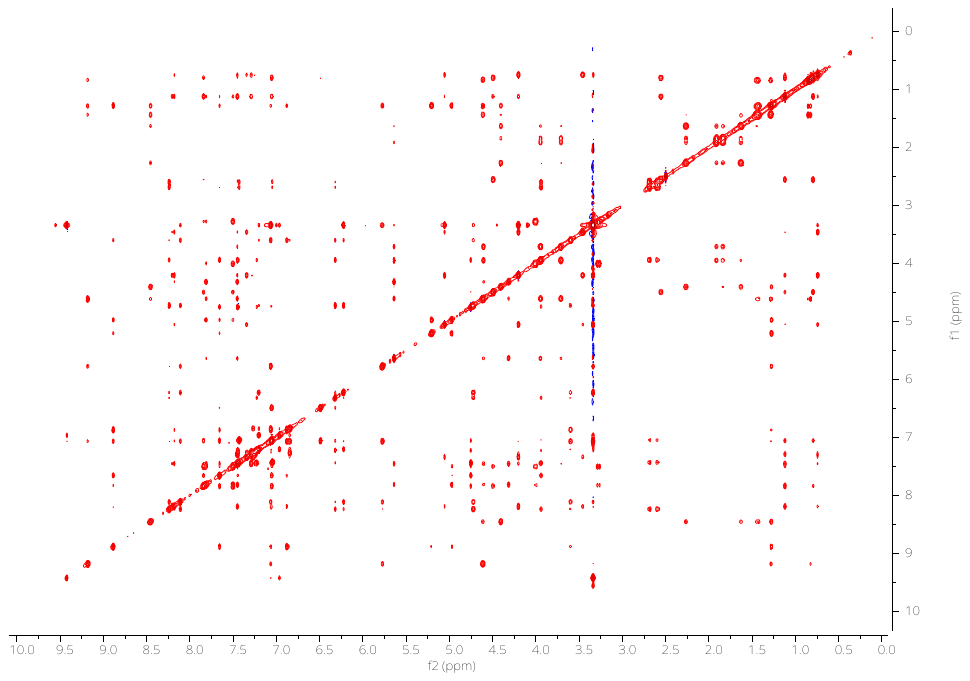

S7 ^1^H-^15^N HSQC Spectrum of Nyuzenamide D (**1**) in DMSO-*d_6_*

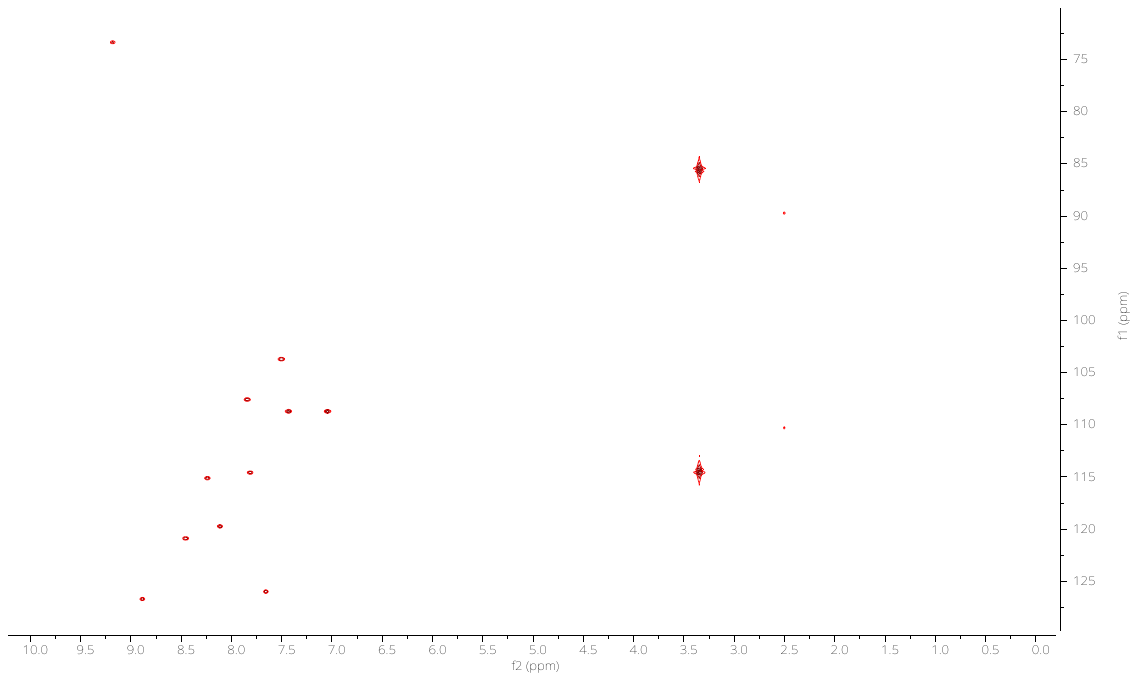

S8 ^1^H-^15^N HMBC Spectrum of Nyuzenamide D (**1**) in DMSO-*d_6_*

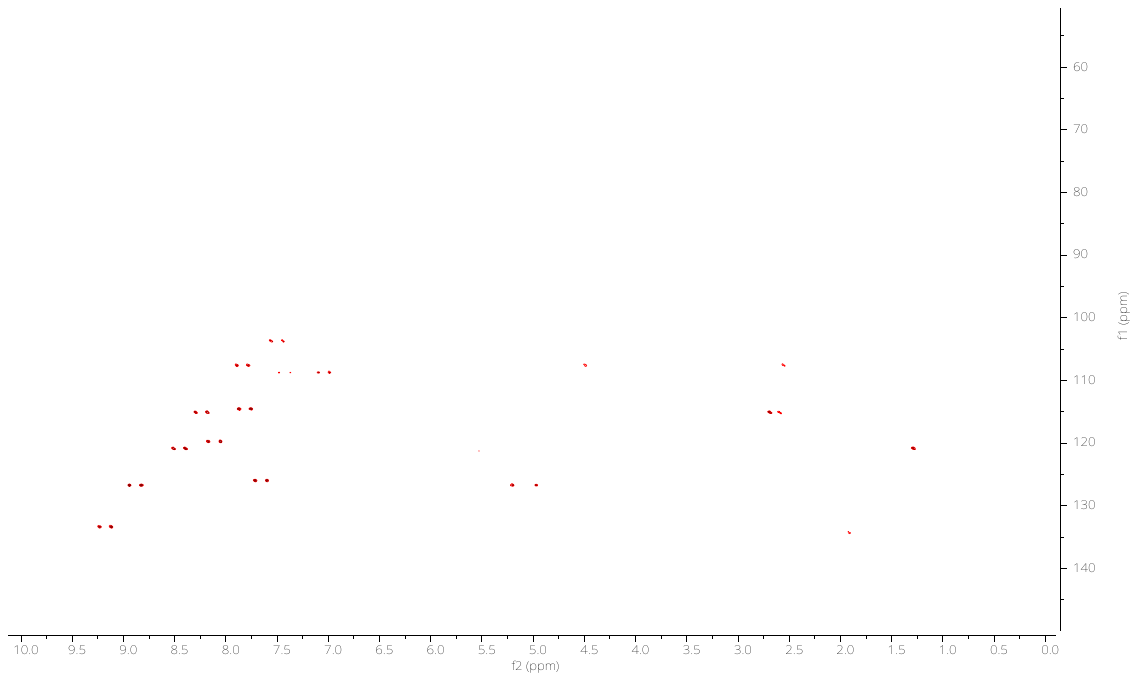

S9 ^1^H NMR (800 MHz) Spectrum of Nyuzenamide E (**2**) in DMSO-*d_6_*

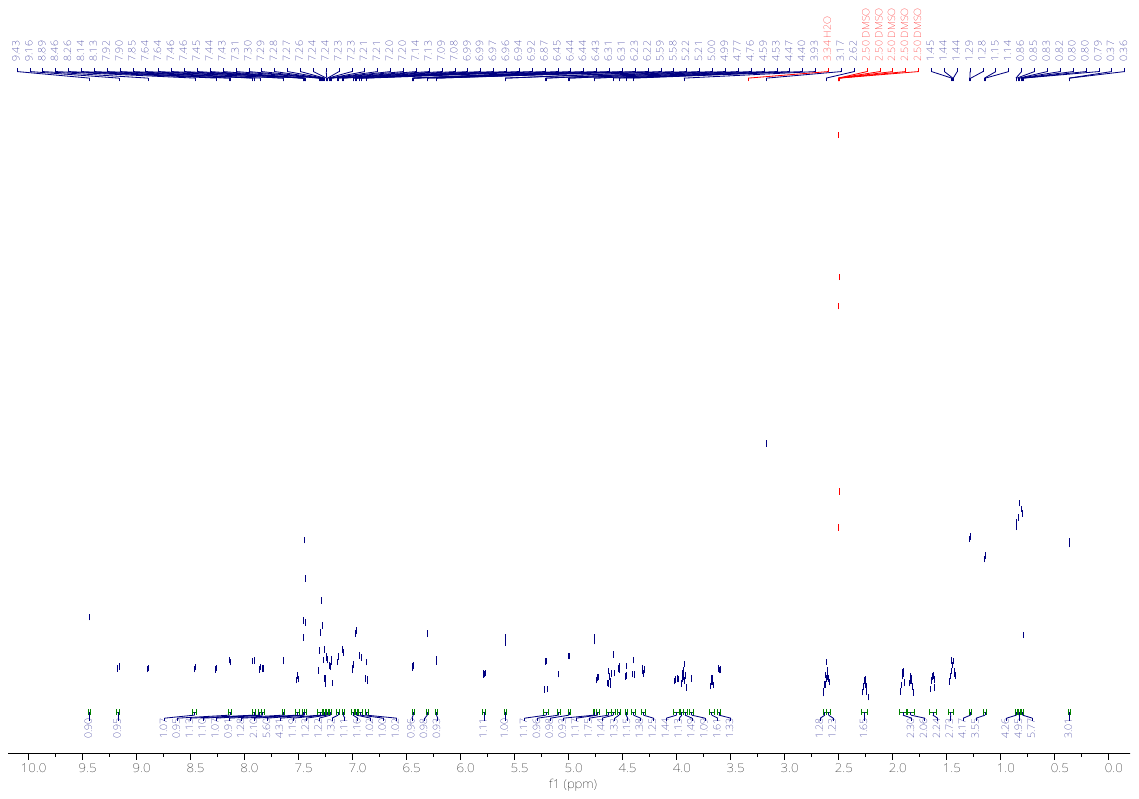

S10 ^13^C NMR (200 MHz) Spectrum of Nyuzenamide E (**2**) in DMSO-*d_6_*

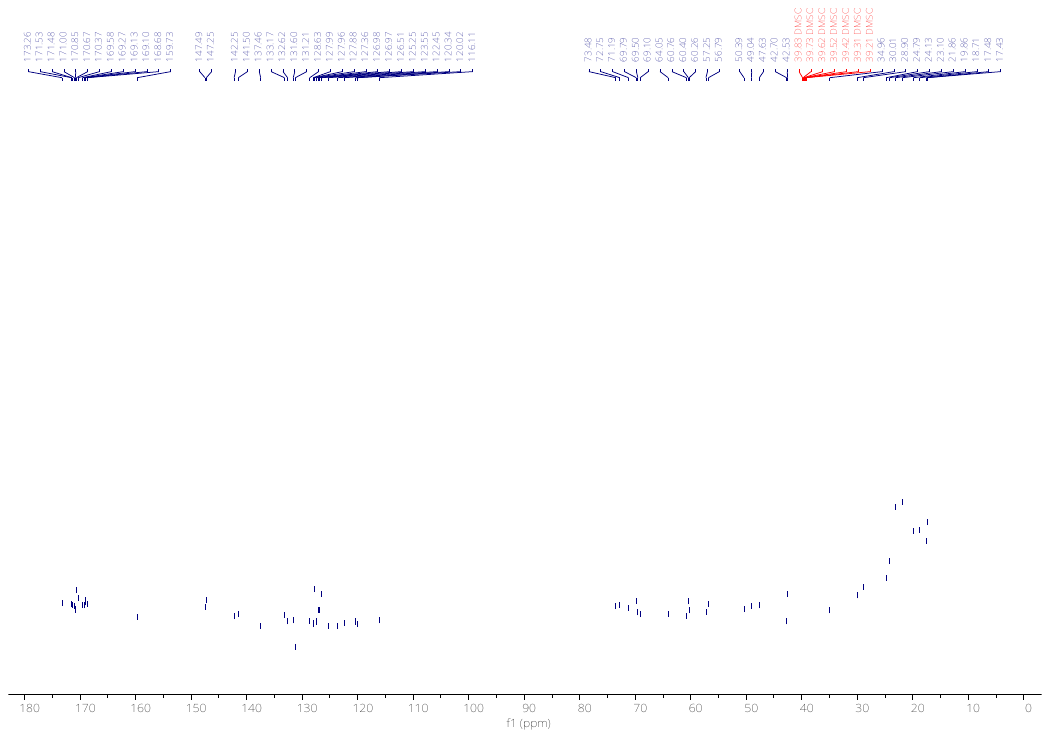

S11 DQF-COSY Spectrum of Nyuzenamide E (**2**) in DMSO-*d_6_*

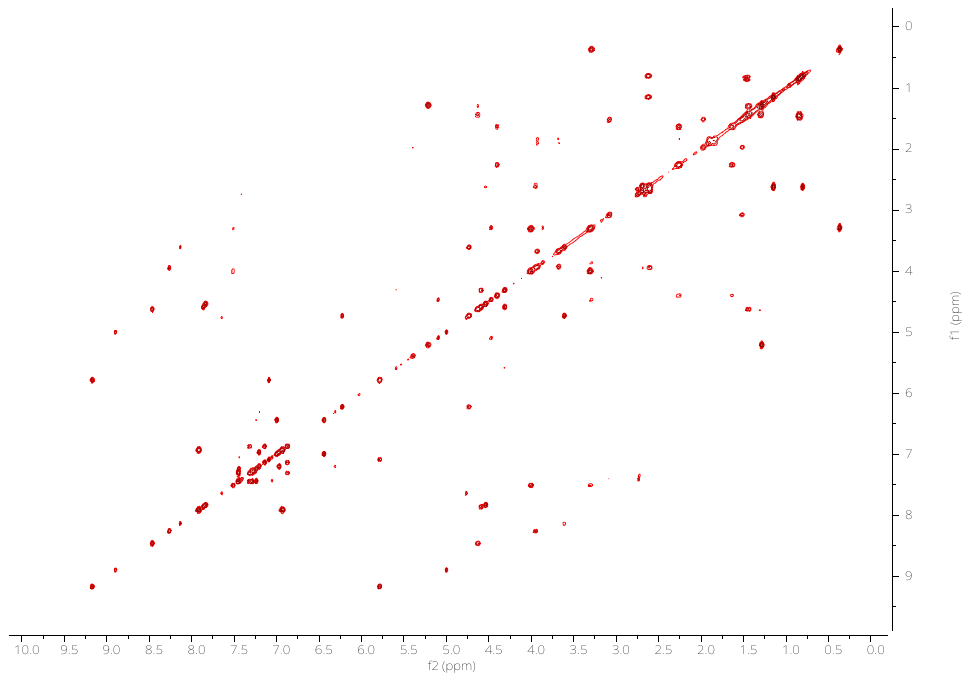

S12 Edited HSQC Spectrum of Nyuzenamide E (**2**) in DMSO-*d_6_*

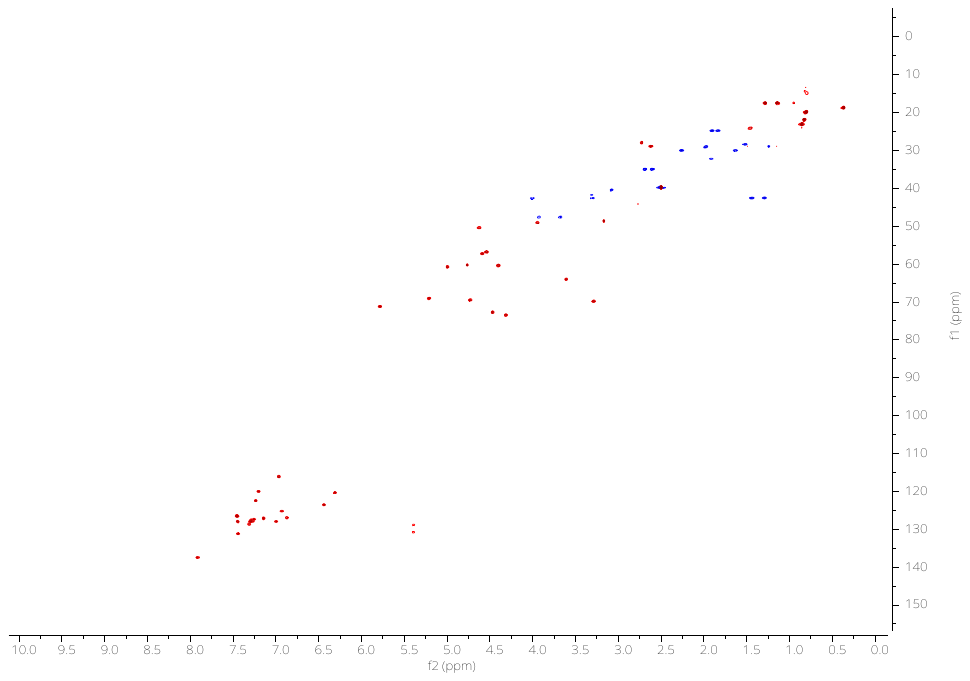

S13 HMBC Spectrum of Nyuzenamide E (**2**) in DMSO-*d_6_*

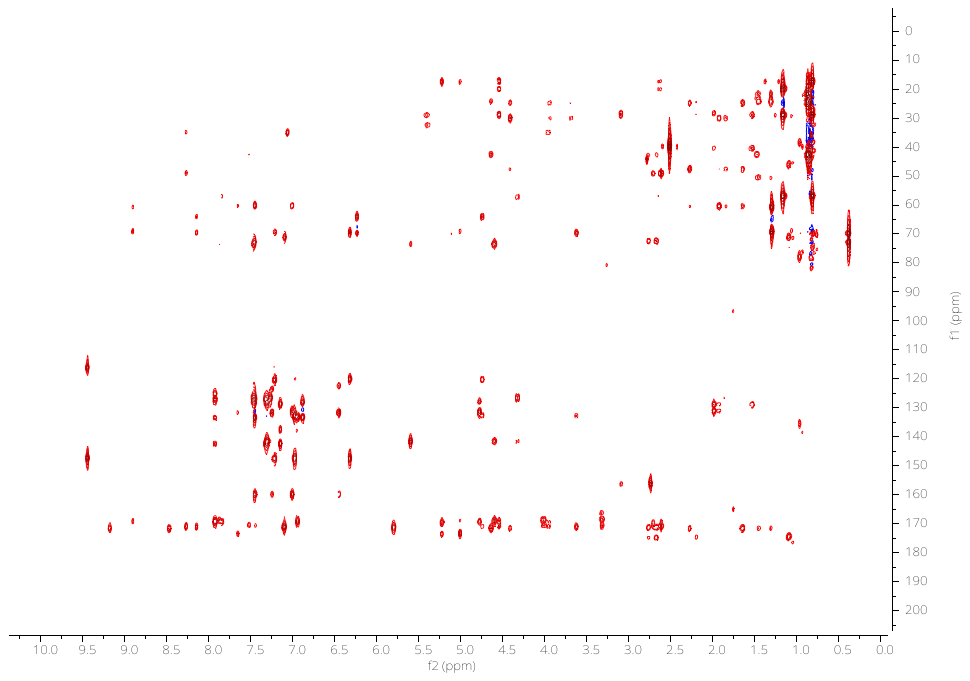

S14 NOESY (600 ms) Spectrum of Nyuzenamide E (**2**) in DMSO-*d_6_*

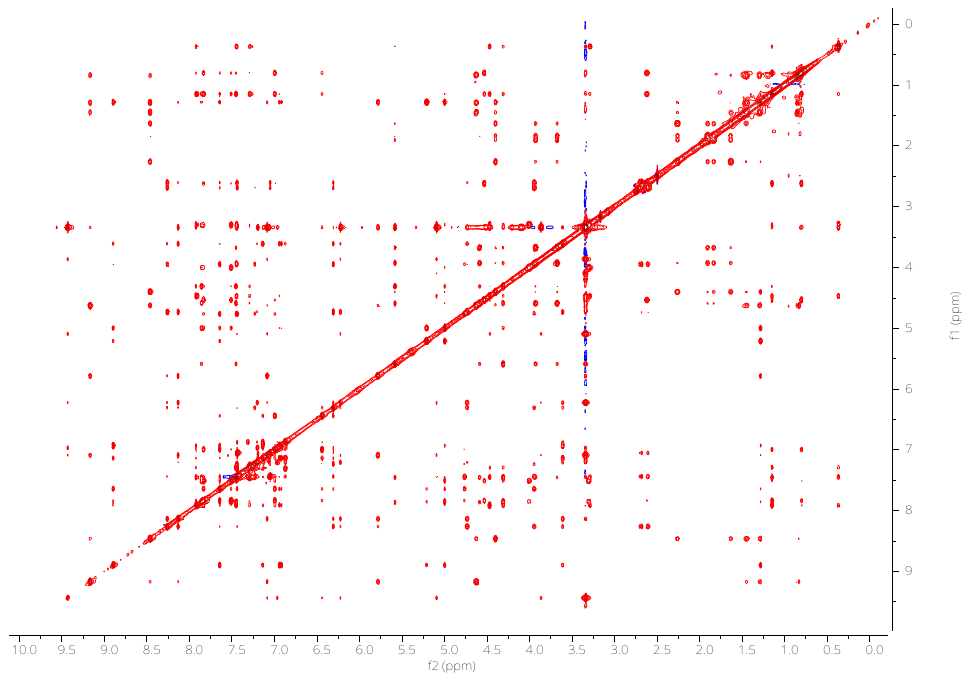

S15 UV Spectrum of Nyuzenamide D (**1**)

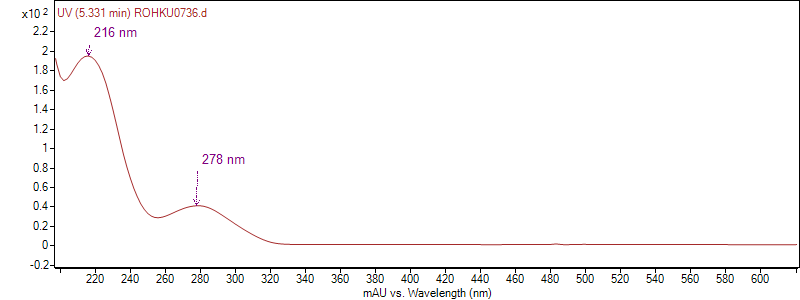

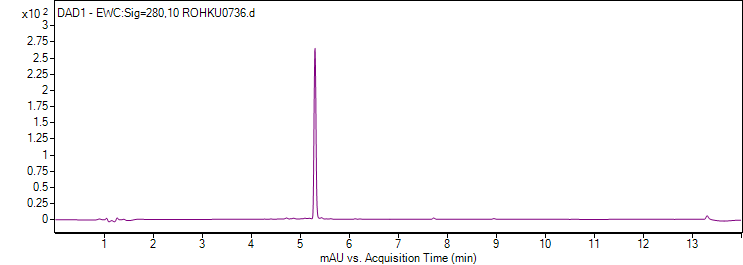

S16 (+)-HRESIMS Spectrum of Nyuzenamide D (**1**)

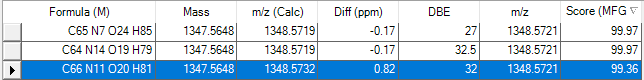

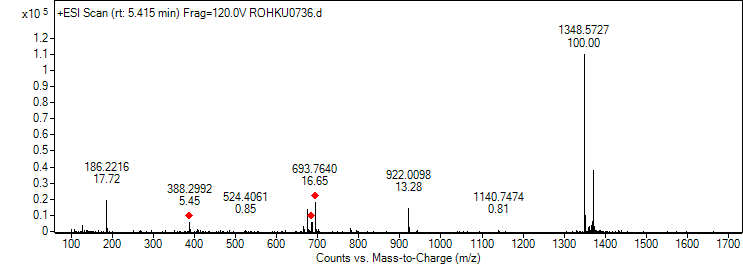

S17 UV Spectrum of Nyuzenamide E (**2**)

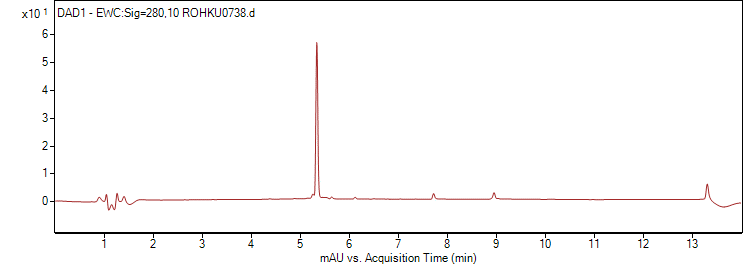

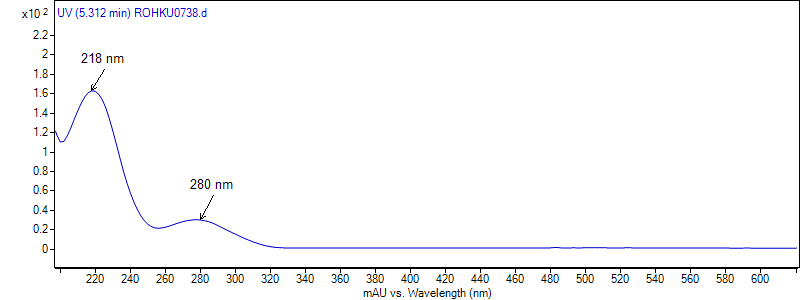

S18 (+)-HRESIMS Spectrum of Nyuzenamide E (**2**)

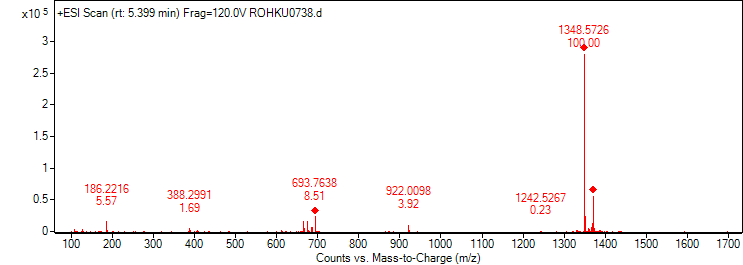

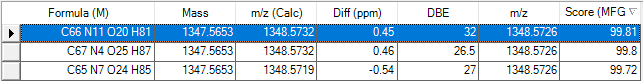

S19 The BGC annotation of *S. hygroscopicus* subsp. *hygroscopicus* DSM 40578.

| **Region** | **Type** | **From** | **To** | **Most similar known cluster** |  | **Similarity** |
| --- | --- | --- | --- | --- | --- | --- |
| Region 1.1 | 2dos | 7,476 | 38,841 | hygromycin A | Saccharide | 69% |
| Region 1.2 | NRP-metallophore, NRPS | 106,202 | 164,544 | coelichelin | NRP | 100% |
| Region 1.3 | terpene | 629,525 | 655,317 | isorenieratene | Terpene | 85% |
| Region 1.4 | NRPS-like | 666,552 | 708,504 | lasalocid | Polyketide | 3% |
| Region 1.5 | T1PKS | 767,974 | 915,446 | niphimycins C-E | Polyketide | 45% |
| Region 1.6 | terpene | 1,022,751 | 1,044,085 | 2-methylisoborneol | Terpene | 100% |
| Region 1.7 | T1PKS, NRPS | 1,055,565 | 1,264,616 | nigericin | Polyketide:Modular type I polyketide | 100% |
| Region 1.8 | hserlactone | 1,486,050 | 1,506,811 |  |  |  |
| Region 1.9 | butyrolactone | 1,511,388 | 1,522,323 |  |  |  |
| Region 1.10 | NAPAA | 1,924,713 | 1,958,849 | griseusin | Polyketide | 5% |
| Region 1.11 | lanthipeptide-class-i | 1,962,243 | 1,986,684 | steffimycin D | Polyketide:Type II polyketide+Saccharide: Hybrid/tailoring saccharide | 8% |
| Region 1.12 | terpene | 2,187,372 | 2,213,822 | hopene | Terpene | 76% |
| Region 1.13 | T2PKS | 2,344,953 | 2,417,468 | spore pigment | Polyketide | 83% |
| Region 1.14 | redox-cofactor | 2,479,222 | 2,501,923 | zorbamycin | NRP:Glycopeptide+Polyketide:Modular type I polyketide+Saccharide: Hybrid/tailoring saccharide | 4% |
| Region 1.15 | RiPP-like | 2,633,542 | 2,644,849 |  |  |  |
| Region 1.16 | ectoine | 2,737,902 | 2,748,330 | ectoine | Other | 100% |
| Region 1.17 | NI-siderophore | 2,828,541 | 2,858,337 | kinamycin | Polyketide | 13% |
| Region 1.18 | NRPS-like | 3,410,650 | 3,453,496 | echoside A/echoside B/echoside C/echoside D/echoside E | NRP | 88% |
| Region 1.19 | triceptide | 3,768,985 | 3,790,350 |  |  |  |
| Region 1.20 | NI-siderophore | 3,976,704 | 4,006,512 | legonoxamine A/desferrioxamine B/legonoxamine B | Other | 100% |
| Region 1.21 | terpene | 5,004,736 | 5,027,141 | geosmin | Terpene | 100% |
| Region 1.22 | arylpolyene, NRPS, lassopeptide | 5,261,263 | 5,372,021 | skyllamycin D/skyllamycin E | NRP | 34% |
| Region 1.23 | indole | 5,677,395 | 5,698,540 | 5-isoprenylindole-3-carboxylate β-D-glycosyl ester | Other | 61% |
| Region 1.24 | NRPS | 5,891,992 | 5,936,245 | ochronotic pigment | Other | 75% |
| Region 1.25 | HR-T2PKS | 5,987,012 | 6,037,306 | cinnapeptin | NRP | 50% |
| Region 1.26 | RiPP-like | 6,045,037 | 6,057,508 | granaticin | Polyketide:Type II polyketide | 10% |
| Region 1.27 | T1PKS | 6,200,450 | 6,386,638 | desulfoclethramycin/clethramycin | Other | 48% |
| Region 1.28 | lassopeptide | 6,836,088 | 6,857,609 | alchivemycin A/alchivemycin B | Polyketide+NRP | 11% |
| Region 1.29 | ectoine | 7,397,781 | 7,408,185 | ectoine | Other | 100% |
| Region 1.30 | NI-siderophore | 7,559,931 | 7,592,202 | peucechelin | NRP | 20% |
| Region 1.31 | T1PKS, amglyccycl | 7,813,465 | 7,945,814 | validamycin A | Saccharide+Other:Cyclitol | 37% |
| Region 1.32 | T1PKS | 8,065,873 | 8,142,196 | butyrolactol A | Polyketide | 80% |
| Region 1.33 | T1PKS, NRPS-like | 8,240,810 | 8,323,276 | geldanamycin | Polyketide | 65% |
| Region 1.34 | terpene | 8,407,049 | 8,428,227 | pristinol | Terpene | 100% |
| Region 1.35 | T1PKS, NRPS-like, NRPS | 8,642,212 | 8,699,644 | hygrocin A/hygrocin B | Polyketide | 41% |
| Region 1.36 | NRPS-like | 8,707,991 | 8,749,850 | aristeromycin | Other | 12% |
| Region 1.37 | RiPP-like | 8,832,072 | 8,842,290 | hexacosalactone A | Other | 9% |
| Region 1.38 | NRPS, lanthipeptide-class-i | 8,846,600 | 8,934,836 | cyclofaulknamycin | Polyketide | 8% |
| Region 1.39 | hydrogen-cyanide | 9,107,160 | 9,120,039 | aborycin | RiPP | 21% |
| Region 1.40 | T1PKS | 9,197,635 | 9,244,015 | streptovaricin | Polyketide | 4% |
| Region 1.41 | terpene | 9,614,927 | 9,635,946 |  |  |  |
| Region 1.42 | 2dos | 9,647,502 | 9,678,867 | hygromycin A | Saccharide | 69% |

S20 Annotation of biosynthetic genes of nyuzenamides D–E in *S. hygroscopicus* subsp. *hygroscopicus* DSM 40578.

| **ORF** | **Size^n^** | **Size^a^** | **Proposed function** | **SI/ID*** | **Protein homologue and origin** |
| --- | --- | --- | --- | --- | --- |
| Orf1 | 816 | 271 | regulatory protein LuxR | 95/92 | AEM88114.1, *Streptomyces violaceusniger* Tu 4113 |
| Orf2 | 810 | 269 | LuxR family transcriptional regulator | 97/95 | WP_161560875.1, *Streptomyces antimycoticus* |
| Orf3 | 744 | 247 | thioesterase | 95/94 | GDY43645.1, *Streptomyces antimycoticus* |
| Orf4 | 729 | 242 | IclR family transcriptional regulator | 95/93 | RSS49760.1, *Streptomyces* sp. WAC05858 |
| Orf5 | 810 | 269 | AfsR/SARP family transcriptional regulator | 100/100 | WP_107449811.1, *Streptomyces* sp. NBRC 109436 |
| Orf6 | 270 | 89 | hypothetical protein | 89/85 | WP_209468855.1, *Streptomyces iranensis* |
| Orf7 | 498 | 165 | hypothetical protein | 97/96 | MBD3009861.1, *Streptomyces* sp. 5-10 |
| NyuA | 252 | 83 | acyl carrier protein | 98/99 | WP_020870443.1, *Streptomyces rapamycinicus* |
| NyuB | 1242 | 413 | beta-ketoacyl-[acyl-carrier-protein] synthase family protein | 98/97 | WP_216347065.1, *Streptomyces* sp. 4503 |
| NyuC | 1134 | 377 | 3-oxoacyl-ACP synthase | 99/99 | WP_064456566.1, *Streptomyces* sp. NBRC 109436 |
| NyuD | 969 | 322 | 3-oxoacyl-ACP synthase | 97/98 | WP_064456567.1, *Streptomyces* sp. NBRC 109436 |
| Orf8 | 1065 | 354 | alpha/beta hydrolase | 94/92 | WP_137966309.1, *Streptomyces antimycoticus* |
| Orf9 | 1020 | 339 | hypothetical protein | 97/96 | MBI0374821.1, *Streptomyces albiflaviniger* |
| NyuE | 1164 | 387 | 3-oxoacyl-ACP synthase | 98//98 | WP_064456569.1, *Streptomyces* sp. NBRC 109436 |
| Orf10 | 516 | 171 | dehydratase | 88/86 | KUL65536.1, *Streptomyces violaceusniger* |
| NyuF | 495 | 164 | beta-hydroxyacyl-ACP dehydratase | 99/99 | WP_064456570.1, *Streptomyces* sp. NBRC 109436 |
| NyuG | 747 | 248 | 3-oxoacyl-[acyl-carrier-protein] reductase | 100/99 | WP_064456571.1, *Streptomyces* sp. NBRC 109436 |
| NyuH | 741 | 246 | 2-hydroxychromene-2-carboxylate isomerase | 96/96 | WP_253371335.1, *Streptomyces* sp. RKCA744 |
| Orf11 | 207 | 68 | MbtH family protein | 97/96 | WP_216344772.1, *Streptomyces* sp. 4503 |
| Orf12 | 2088 | 695 | AarF/UbiB family protein | 99/99 | WP_064456573.1, *Streptomyces* sp. NBRC 109436 |
| NyuI | 1110 | 369 | prephenate dehydrogenase | 99/99 | WP_064456576.1, *Streptomyces* sp. NBRC 109436 |
| NyuJ | 2784 | 927 | Aminotransferase class I/II-fold pyridoxal phosphate-dependent enzyme | 92/92 | WP_079150261.1, *Streptomyces* sp. NBRC 109436 |
| NyuK | 1044 | 347 | 4-hydroxyphenylpyruvate dioxygenase | 94/92 | WP_020870456.1, *Streptomyces rapamycinicus* |
| Orf13 | 684 | 227 | Hypothetical protein | 95/90 | EXU63499.1, *Streptomyces* sp. PRh5 |
| NyuL | 1227 | 408 | Cytochrome P450 | 99/99 | WP_064456577.1, *Streptomyces* sp. NBRC 109436 |
| Orf14 | 975 | 324 | ATP-binding cassette domain-containing protein | 98/96 | WP_191069122.1, *Streptomyces* sp. 5-10 |
| Orf15 | 795 | 264 | ABC transporter permease | 98/95 | WP_044573534.1, *Streptomyces iranensis* |
| Orf16 | 300 | 99 | Hypothetical protein | 82/73 | GHH26059.1, *Streptomyces rubradiris* |
| Orf17 | 1011 | 336 | Methyltransferase domain-containing protein | 92/87 | WP_223124033.1, *Streptomyces cacaoi* |
| Orf18 | 432 | 143 | lasso peptide biosynthesis B2 protein | 92/91 | WP_063793421.1, *Streptomyces hirsutus* |
| Orf19 | 303 | 100 | PqqD family peptide modification chaperone | 87/84 | WP_149563218.1, *Streptomyces cacaoi* |
| Orf20 | 1773 | 590 | albusnodin/ikarugamycin family macrolactam cyclase | 91/87 | WP_152262678.1, *Streptomyces mobaraensis* |
| Orf21 | 126 | 41 | Hypothetical protein | 100/100 | WP_284709075.1, *Streptomyces* sp. |
| Orf22 | 1275 | 424 | helix-turn-helix domain-containing protein | 93/89 | WP_229852203.1, *Streptomyces albospinus* |
| NyuM | 1074 | 357 | LLM class flavin-dependent oxidoreductase | 97/93 | WP_044573537.1, *Streptomyces iranensis* |
| NyuN | 1218 | 405 | Cytochrome P450 | 98/96 | MBA6439831.1, *Streptomyces* sp. GMR22 |
| NyuO | 11361 | 3786 | Non-ribosomal peptide synthetase (CAT-CAT-CATE) | 97/97 | WP_064456585.1, *Streptomyces* sp. NBRC 109436 |
| NyuP | 11970 | 3989 | Non-ribosomal peptide synthetase (CAT-CATE-CAT-Te) | 87/83 | WP_164425344.1, *Streptomyces rhizosphaericus* |
| NyuQ | 1260 | 419 | Cytochrome P450 | 98/95 | WP_020870465.1, *Streptomyces rapamycinicus* |
| NyuR | 477 | 158 | Beta-hydroxyacyl-ACP dehydratase | 97/96 | WP_107449898.1, *Streptomyces* sp. NBRC 109436 |
| Orf23 | 375 | 124 | Hypothetical protein | 91/88 | WP_198278730.1, *Streptomyces javensis* |
| NyuS | 15219 | 5072 | Non-ribosomal peptide synthetase (CAT-CAT-CAT-CATE) | 95/95 | WP_064457897.1, *Streptomyces* sp. NBRC 109436 |
| Orf(+1) | 510 | 169 | Flavin reductase family protein | 94/92 | WP_037956334.1, *Streptomyces* sp. PRh5 |
| Orf(+2) | 639 | 212 | MBL fold metallo-hydrolase | 98/95 | WP_208515409.1, *Streptomyces* sp. NEAU-YJ-81 |

n, The size of nucleotides; a, The size of amino acids; *, similarity-identity ratio.

S21 Sequence alignment of the amino acids of peptide-carrier protein domain.

Consensus	EEILCGLFAEVLGL--PEVGVDDSFFDLGGHSLLATRLVXRARAALGVELSVRDVFEAPTVAALAALLDGA 69

EJMJBBLN_45180_PCP.1 EEILCALFAELLGV--PEVGVDDDFFELGGHSLLATRLVTRTRAIFGVELSLRAVFETPSPAAIARGLDRA 69

EJMJBBLN_45180_PCP.2 EAALCALFAEVLGV--PAAGVDDNFFDLGGHSLLATRLTSRIRTTLGAELSVRDLFESPTPAGLAGLVG-- 67

EJMJBBLN_45180_PCP.3 EETLRRLVAEVLGIGADEVGVDDSFFSLGGDSIVSIQLVARARAA-GLVFTPKDVFRHQTVAALAAAA--- 67

EJMJBBLN_45190_PCP.1 EEILCGLFMETLGL--NAVGIHDDFFELGGHSLHATRLVSRARSAFAVELPLRVLWEAPTVARLAARLRG- 68

EJMJBBLN_45190_PCP.2 EELLCSLFAEVLGL--ERIGVHDSFFDTGGDSIMSIQLVRRARNA-GIKLTPRDVFEQKTVAGLAALTA-- 66

EJMJBBLN_45190_PCP.3 EEILCGLFREELGL--DVVGVHDSFFDLGGDSIMSMSLVNRIRSAFGVKLGMRTVLTAPTVAELALRLGG- 68

EJMJBBLN_45230_PCP.1 EQVLAGLFAEVLGV--PEVGVDDNFFALGGHSLAATRLAGRARSVLGVELAVHEVFDRPTVAALAGLLDT- 68

EJMJBBLN_45230_PCP.2 EQVLCGLFAEILEL--ASVSVDDNFFSLGGHSLLATRLVLRIQAVLGADVAVRDLFDAPTVAELARLVDRA 69

EJMJBBLN_45230_PCP.3 EEILCGLFAELLGV--GPVSVDDDFFDLGGHSLLAARLVNRIRTALDAELSVRTLFEAPTVAALATALDGR 69

EJMJBBLN_45230_PCP.4 EEALCAVFAEVLRL--TSVGIDDSFFELGGDSIVSMQLIARARAA-GLVLRVGDVFEHKTVAGLAAVAE-- 66

S22 The strains and plasmids used in this study.

| **Strains** | **Description** |
| --- | --- |
| One Shot™ Mach1™ T1 Phage-Resistant Chemically Competent *E. coli* | For routine plasmids maintenance and cloning |
| *E. coli* ET12567/pUZ8002 | For conjugating plasmids into *Streptomyces* |
| *S. hygroscopicus* subsp. *hygroscopicus* DSM 40578 | Wild-type strain |
| *S. hygroscopicus* subsp. *hygroscopicus* DSM 40578*/*Δ*nyuS* | Knock-out of Δ*nyuS* in *S. hygroscopicus* subsp. *hygroscopicus* DSM 40578 |
| **Plasmids** |  |
| pCRISPR-Cas9 | Gene knockout/in for actinomycetes |
| pCRISPR-Cas9/Δ*nyuS* | Modified plasmid for knock-out of *nyuS* |

S23 The primers used in this study.

| **Primer name** | **Sequence (5’ → 3’)** | **Description** |
| --- | --- | --- |
| Del-*nyuS*-sgRNA | CCGGTTGGTAGGATCGACGGGagtcgtcgtcgaccgacaccGTTTTAGAGCTAGAAATAGC | Knock-out of *nyuS,* the base marked in red is sgRNA sequence |
| Del-*nyuS*-uparm-F | tcgtcgaaggcactagaaggtgttccttgatccgcttgac | Forward primer for amplification of upstream homologous flank of *nyuS* |
| Del-*nyuS*-uparm-R | gaacacgtcttcctgatcgtgtgatgtgttcgagcacaag | Reverse primer for amplification of upstream homologous flank of *nyuS* |
| Del-*nyuS*-downarm-F | cttgtgctcgaacacatcacacgatcaggaagacgtgttc | Forward primer for amplification of downstream homologous flank of *nyuS* |
| Del-*nyuS*-downarm-R | ggtcgatccccgcatataggccgatgaccaggtgaagatc | Reverse primer for amplification of downstream homologous flank of *nyuS* |
| ID-*nyuS*-F1 | cagcagtccgtagccgatc | Forward primer for screening *nyuS* mutants |
| ID-*nyuS*-R1 | gaacacgtcttcctgatcgt | Reverse primer for screening *nyuS* mutants |
| ID-*nyuS*-F2 | ccgtcttgtgctcgaacac | Forward primer for screening *nyuS* mutants |
| ID-*nyuS*-R2 | ccgagaacgacctggactg | Reverse primer for screening *nyuS* mutants |

S24 Antibacterial and anticandidal activities of **1** and **2**.

| Microorganisms | Minimum Inhibitory Concentration (MIC)(µg/mL) | | | | |
| --- | --- | --- | --- | --- | --- |
|  | **1** | **2** | **Gentamicin** | **Chloramphenicol** | **Amphotericin b** |
| *Bacillus subtilis* | >64 | >64 | <0.25 | 4 | NT |
| *Staphylococcus aureus* | >64 | >64 | 2 | 8 | NT |
| *Pseudomonas aeruginosa* PAO1 | >64 | >64 | 4 | 32 | NT |
| *Candida albicans* IBT654 | >64 | >64 | NT | NT | <0.25 |

NT, not tested.

S25 Antifungal activity of **1** and **2** against (**A**) *Alternatia solani*, (**B**) *Botrytis cinerea*, (**C**) *Aspergillus flavus*, (**D**) *Aspergillus niger*, (**E**) *Aspergillus fumigatus*, (**F**) *Aspergillus tubingensis*.

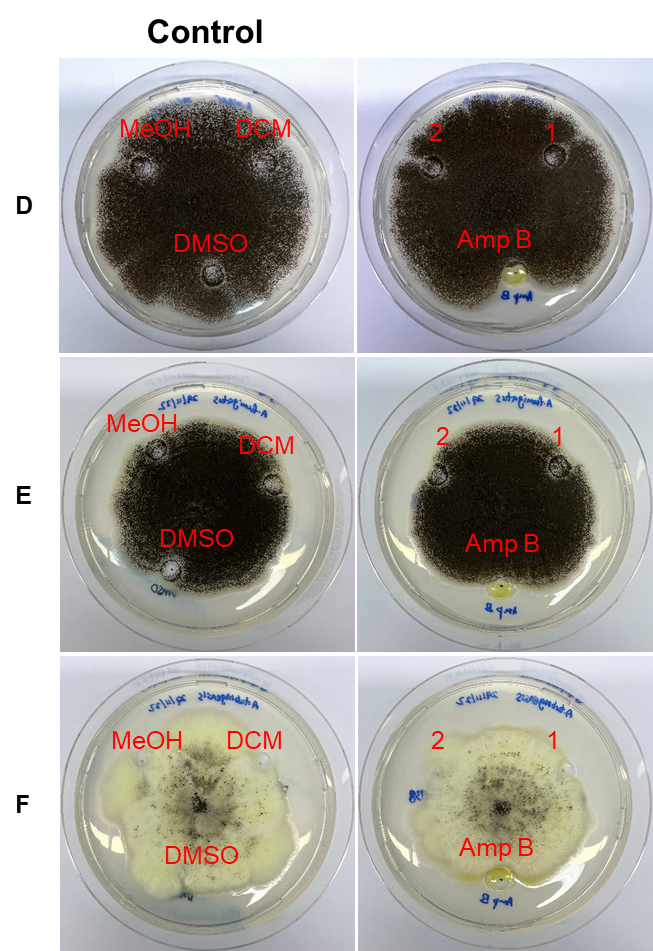

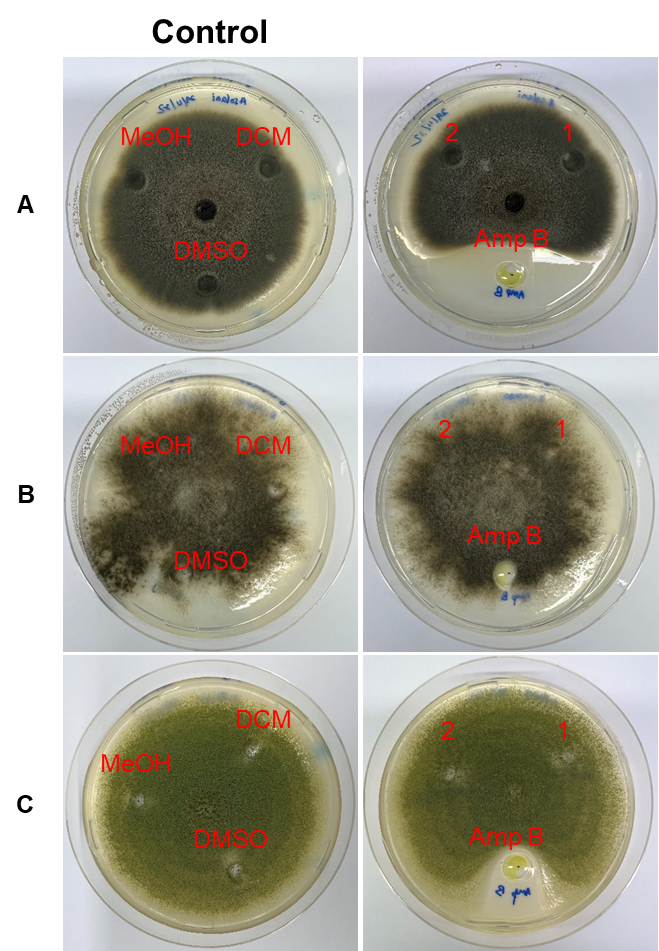
